# Removing direct photocurrent artifacts in optogenetic connectivity mapping data via constrained matrix factorization

**DOI:** 10.1101/2023.07.13.548849

**Authors:** Benjamin Antin, Masato Sadahiro, Marta Gajowa, Marcus A. Triplett, Hillel Adesnik, Liam Paninski

**Affiliations:** Mortimer B. Zuckerman Mind Brain Behavior Institute, Columbia University, NY; Grossman Center for the Statistics of Mind, Columbia University, NY; Center for Theoretical Neuroscience, Columbia University, NY; Department of Statistics, Columbia University, NY; Department of Molecular and Cell Biology, University of California, Berkeley, CA

## Abstract

Monosynaptic connectivity mapping is crucial for building circuit-level models of neural computation. Two-photon optogenetic stimulation, when combined with whole-cell recordings, has the potential to map monosynaptic connectivity at an unprecedented scale. However, optogenetic mapping of nearby connections poses a challenge, due to stimulation artifacts. When the postsynaptic cell expresses opsin, optical excitation can directly induce current in the patched cell, confounding connectivity measurements. This problem is most severe in nearby cell pairs, where synaptic connectivity is often strongest. To overcome this problem, we developed a computational tool, Photocurrent Removal with Constraints (PhoRC). Our method is based on a constrained matrix factorization model which leverages the fact that photocurrent kinetics are consistent across repeated stimulations at similar laser power. We demonstrate on real and simulated data that PhoRC consistently removes photocurrents while preserving synaptic currents, despite variations in photocurrent kinetics across datasets. Our method allows the discovery of synaptic connections which would have been otherwise obscured by photocurrent artifacts, and may thus reveal a more complete picture of synaptic connectivity. PhoRC runs faster than real time and is available at https://github.com/bantin/PhoRC.

## 1 Introduction

Monosynaptic connectivity mapping provides essential information for constraining models of neural circuits. In visual cortex, for instance, knowledge of spatial connectivity statistics between cells offers important constraints when designing circuit models of perception [Huang et al., 2019, Palmigiano et al., 2023]. Two photon optogenetics, combined with whole-cell patch clamp electrophysiology, allows the mapping of monosynaptic connectivity at a throughput not attainable with traditional paired patch electrophysiology [Packer et al., 2012, Baker et al., 2016, Hage et al., 2022, McRaven et al., 2020, Triplett et al., 2022]. In this approach, a postsynaptic neuron is recorded intracellularly while individual presynaptic neurons are excited by optogenetic stimulation. Recent computational work has shown that this approach can be scaled to test hundreds of potential connections onto a single postsynaptic partner at high speed to obtain a comprehensive spatial map of synaptic connectivity [Draelos et al., 2020, Triplett et al., 2022].

Ideally, opsin would be densely expressed in the cell population to allow thorough probing of the putative presynaptic population. This can yield an “all-to-one” description of connectivity onto a single postsynaptic neuron. However, if both the presynaptic population and the postsynaptic cell express opsin, photostimulation near or directly on the postsynaptic cell can result in recording photocurrents, which confound synaptic currents and connectivity estimation. This artifact has been a frequent practical concern, especially when mapping connections between excitatory neurons [Shemesh et al., 2017, Hage et al., 2022, Printz et al., 2021].

Previously, practitioners adjusted the parameters of the experimental preparation to minimize the effect of this artifact. For example, when opsin expression is sparsely limited across the cell population, then it becomes possible to patch onto an opsin negative cell and avoid the issue altogether. However, sparse expression limits the number of probable connections and limits each experiment to only being able to acquire partial connectivity maps. Thus in a previous study, practitioners aggregated partial map data across multiple experiments from different patched cells [Hage et al., 2022].

Advances in optogenetic tools, including opsins [Baker et al., 2016, Shemesh et al., 2017] and two-photon optics [Pégard et al., 2017], have improved upon the effective spatial resolution of optogenetic stimulation. However, direct photocurrent artifacts still pose a major challenge when stimulating presynaptic neurons near the postsynaptic cell [Hage et al., 2022, Printz et al., 2021]. Importantly, it is precisely in the region immediately surrounding a patched cell that we expect connectivity to be strongest for many cell types [Holmgren et al., 2003, Hage et al., 2022]. Furthermore, the precise spatial statistics of connectivity are a key parameter in theories of circuit function [Huang et al., 2019, Palmigiano et al., 2023]. It is therefore critical to avoid systematic biases due to opsin contamination when estimating these circuit parameters.

Prior attempts to computationally remove photocurrents have relied on extensive manual curation [Printz et al., 2021], or assumed exact *a priori* knowledge of photocurrent kinetics [Merel et al., 2016] (though see [Triplett et al., 2022] for an exception, discussed in more detail below). The ability to automatically detect and remove photocurrents while preserving underlying postsynaptic currents would enable more complete mapping of local connectivity without limiting the location or number of presynaptic targets. This would allow for the construction of more detailed and accurate circuit models of neural computation.

Towards these aims, we developed and validated a constrained matrix-factorization model which we call Photocurrent Removal with Constraints (PhoRC). Unlike prior approaches, PhoRC removes direct photostimulation artifacts and preserves recorded postsynaptic currents without prior knowledge of the photocurrent waveform. This means it can adapt seamlessly to variability between datasets, or even between different opsins. We validated PhoRC on simulated and real excitatory-to-excitatory mapping data, and found that it enabled consistent removal of photocurrent artifacts and thereby improved improved subsequent connectivity estimation.

## 2. Results

### 2.1 Two mapping protocols demonstrate the confounding effect of direct photocurrents

We consider synaptic connectivity mapping experiments in which currents are recorded from an opsin-expressing postsynaptic cell, and presynaptic targets are optogenetically stimulated with a laser. In this setup, stimulation evokes direct photocurrents in the patched cell whenever the laser is used to stimulate targets that are close to the patched cell (Figure 1a, left). Direct photocurrent can be distinguished from synaptic current by its kinetics: immediate onset when stimulation begins, followed by roughly exponential decay when stimulation ends [Rickgauer and Tank, 2009, Schoeters et al., 2021, Mardinly et al., 2018, Sridharan et al., 2022]. Laser power is expected to modulate the amplitude of photocurrent response, with little impact on latency or overall kinetics [Rickgauer and Tank, 2009, Packer et al., 2012]. By contrast, typical PSCs have longer and more variable latencies compared to photocurrents, with different rise times (Figure 1a).

**Figure 1:**
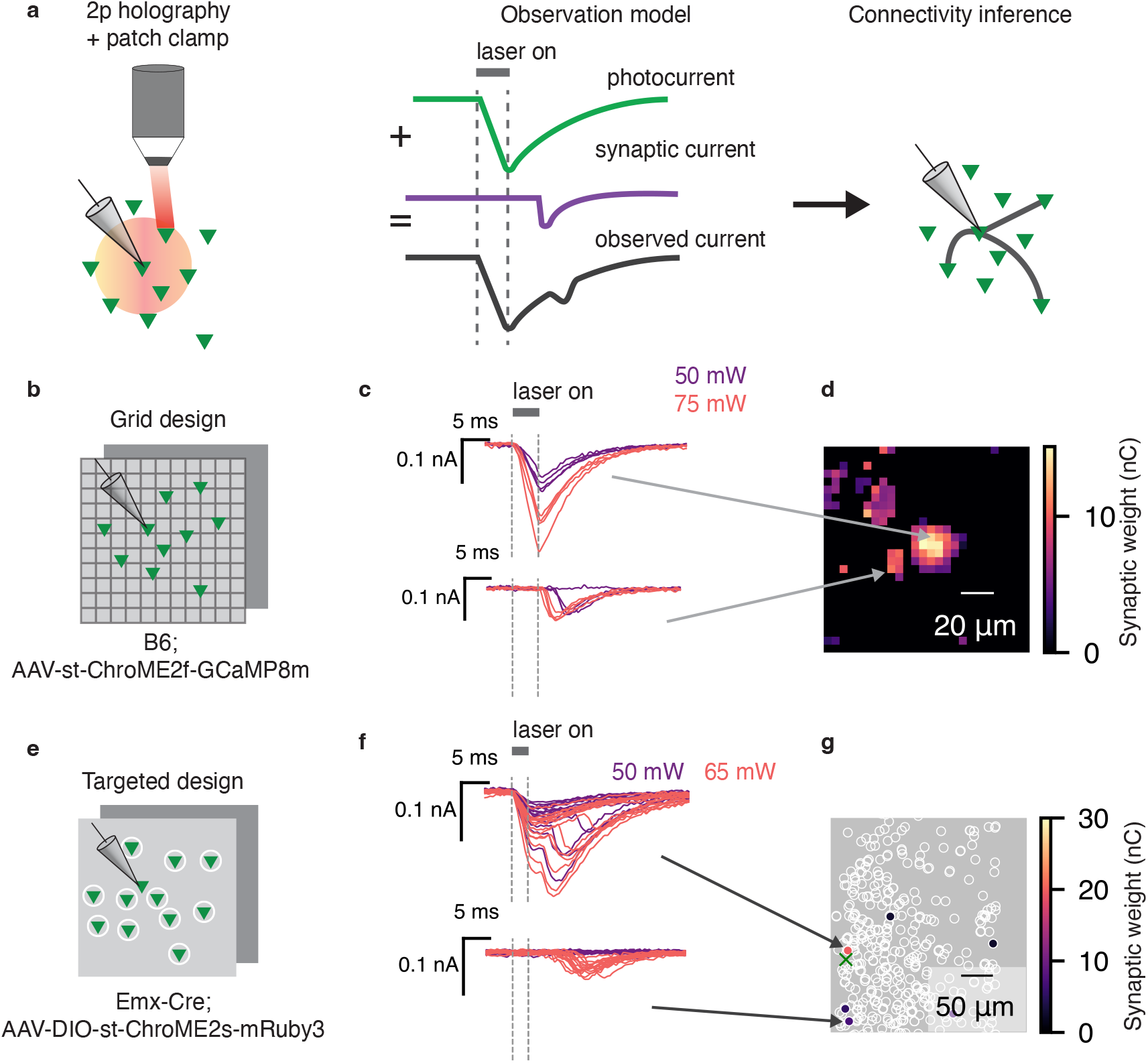
Photocurrent contamination confounds mapping of nearby synaptic connections. **a**, Synaptic connectivity mapping is made possible by patching a single downstream cell and targeting surrounding cells with two-photon excitation. When the downstream cell expresses opsin, the observed currents contain a mix of synaptic current and direct photocurrent. Left panel: cartoon of experimental setup; red circle represents radius in which direct photocurrent will be evoked. Middle panel: photocurrent artifacts are added to any observed synaptic current signals. Right panel: such artifacts subsequently corrupt connectivity inference. **b**, Schematic of grid experimental design. Stimulation laser was fired randomly in a grid pattern to excite cells expressing ChroME2f. **c**, Representative traces from a proximal (top) and distal (bottom) stimulus location in a grid experiment. Without accounting for the photocurrent artifact, both locations are naively categorized as being connected to the patched cell. The top traces represent putative photocurrent-only responses: immediate rise while the laser is on, with amplitude modulated by laser power. The bottom traces represent prototypical EPSC responses. Note the steeper rise, with onset a few milliseconds after stimulation. EPSC response latency is modulated by laser power. Purple and red traces denote 30 and 50 mW stimulation, respectively. **d**, Inferred connectivity map for an example grid experiment. The cluster of strong inferred weights in the center is likely artifactual. *Caption continued on next page*. **e**, In the targeted design, excitatory cells expressed ChroME2s fused to mRuby3, allowing us to detect and segment presynaptic target cells. **f**, Representative traces from a proximal (top) and distal (bottom) target cell in a targeted experiment. The top traces contain a putative mixture of photocurrent and synaptic current. Bottom traces represent putative EPSC-only response. **g**, Inferred connectivity from a representative targeted experiment. Green cross denotes location of the postsynaptic cell.

When the postsynaptic cell is directly photoactivated, incoming synaptic currents will be added to the induced photocurrent, creating a corrupted measurement, and subsequently biasing connectivity inference. Thus for each location stimulated, there are several possible scenarios: we may evoke pure photocurrent, pure synaptic current, or a mix of the two (Figure 1a). An effective tool for photocurrent subtraction would address all three of these cases. We therefore asked whether we could design a computational tool capable of subtracting photocurrents while preserving synaptic currents, even when they are mixed together.

We employed two mapping protocols in order to systematically study this question (Figure 1). In both sets of experiments we mapped connections between pyramidal cells by combining the scanless computer-generated holographic optogenetic system named 3D-SHOT [Pégard et al., 2017] with voltage-clamp electrophysiology, and using the soma-targeted ChroME family of opsins [Sridharan et al., 2022]. Following similar work, we titrated laser power to almost always evoke zero or one action potentials following brief (3-5ms) pulses of stimulation [Packer et al., 2012, Hage et al., 2022, Triplett et al., 2022]. At each stimulation, the response was taken to be the total charge transfer during a short window following stimulation (see subsection 4.2).

In the first protocol, which we term the grid-based approach, we pan-neuronally expressed soma-targeted ChroME2f, and recorded intracellular currents from an opsin-positive cell. We then stimulated points at random in a three-dimensional grid surrounding the patched cell (Figure 1b). The grid was approximately 170 *µ*m per side, and included five axial planes spaced 25 *µ*m apart (see section 4.2.1). The grid-based approach allowed us to carefully characterize how photocurrent activation changes with distance and laser power, and was crucial in developing our photocurrent subtraction method. Using the responses following each stimulation, we inferred connectivity using the CAVIaR software package [Triplett et al., 2022]. In brief, CAVIaR uses a neural-network based denoiser, combined with a statistical model, to infer an effective synaptic weight for each presynaptic target. This yielded a spatial map in which each pixel was assigned an effective synaptic weight (Figure 1d).

We consistently found a region near the patched cell in which photocurrents dominated the observed signal (Figure 1c,d). The size and shape of this region varied significantly between datasets, ranging from 25-70 *µ*m in diameter (Figs. S1-S5). This led us to infer connections near the patched cell which were actually due to photocurrent artifact. By contrast, connections inferred far from the patched cell displayed typical profiles of excitatory postsynaptic currents (EPSCs) (Figure 1c).

We then further investigated the impact of photocurrent artifacts using a second experimental protocol, which we term the targeted protocol. In this protocol, we expressed a slower opsin (ChroME2s) fused to the fluorescent reporter mRuby3 in excitatory neurons. At the beginning of the experiment, two-photon imaging of mRuby3 was used to detect and segment opsin-expressing cells for subsequent stimulation (Figure 1e). As in the prior set of experiments, we observed substantial direct photocurrent when mapping cells adjacent to the patched cell, which subsequently corrupted estimates of connectivity. As in our grid experiments, inferred connections close to the patched cell frequently contained either pure photocurrent, or a mix of synaptic and photocurrent (Figure 1f,g), and further motivated the development of an automated photocurrent removal tool.

### 2.2 A constrained low-rank model separates photocurrents from synaptic currents

We collected four datasets using the targeted approach and two using the grid approach. For each dataset, we rescaled and plotted the ten largest traces, as determined by signal magnitude during stimulation. We found that even when using the same opsin and stimulating at similar laser powers, there was significant variability in the photocurrent waveform across datasets, possibly due to differing experimental parameters such as access resistance or slice temperature (Figure 2a,b). Although practioners can discard cells with high access resistance, even within the range of recordings which are normally acceptable, there can be substantial variation in access resistance, leading to variations in photocurrent and synaptic current kinetics. This variability poses a challenge to previous automated photocurrent subtraction approaches, which assume prior knowledge of the photocurrent waveform [Merel et al., 2016]. We therefore sought to develop a method which would automatically learn the photocurrent waveform from data. As we show below, pursuing such an approach has the added benefit of automatically adapting to the kinetics of different opsins.

**Figure 2:**
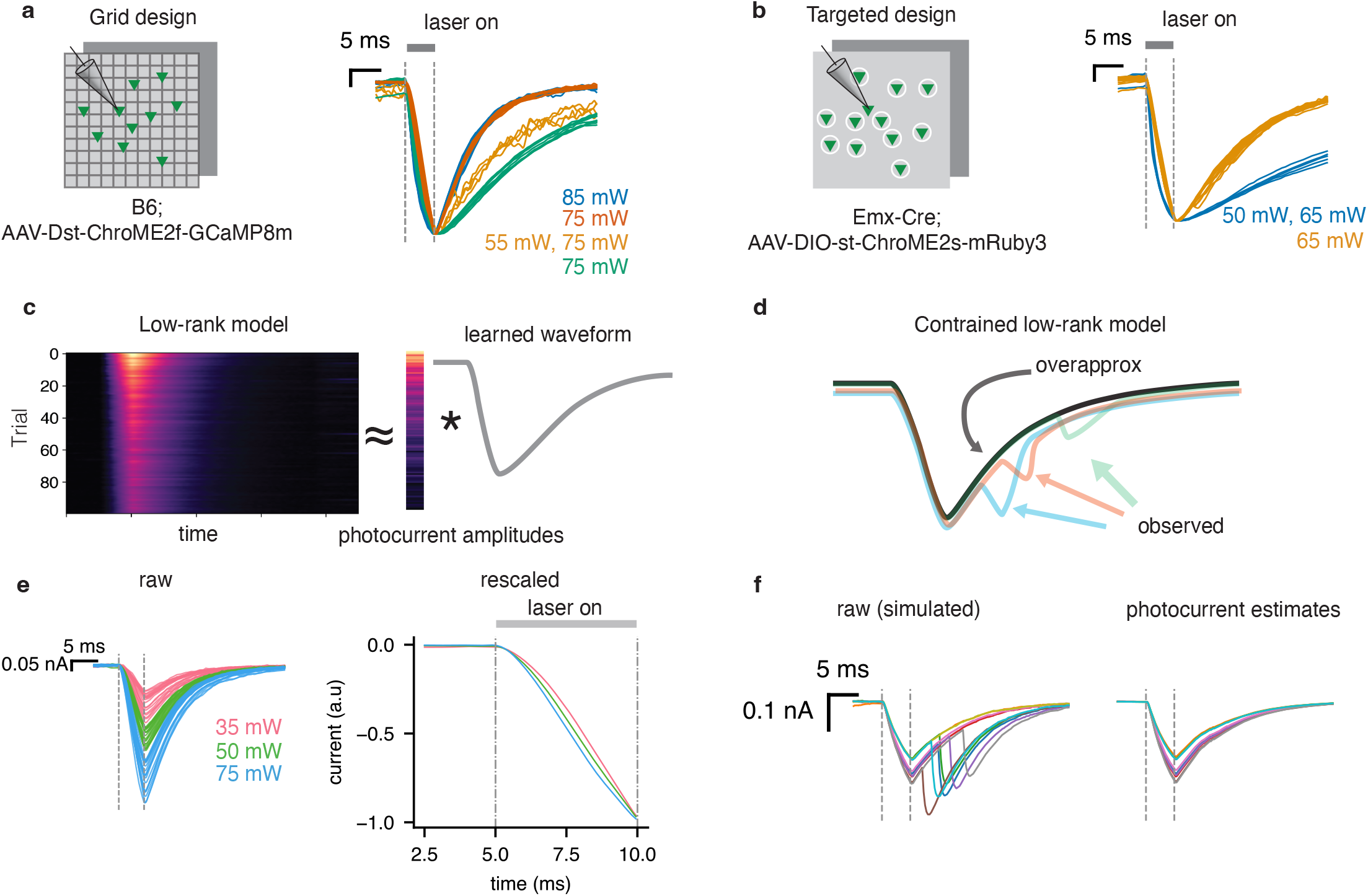
A constrained low-rank model separates synaptic currents from photocurrents. **a**, Scaled photocurrent traces from the grid experiments with ChroME2f. Colors correspond to different datasets; stimulation power used is shown on the right. Even with the same opsin, there is substantial cell-to-cell variability in the photocurrent waveform. **b**, Same as **a**, but for targeted experiments using ChroME2s. **c**, Schematic of the low-rank approach. The matrix of PSC traces (left) can be approximated as a single waveform scaled by a vector of amplitudes. **d**, The overapproximation constraint stops the estimated waveform from picking up PSCs. **e**, Left panel: representative photocurrent traces from a single ChroME2f experiment at three laser powers. Photocurrents scale with laser power. Right panel: For each laser power, we averaged the traces shown in the left panel, and scaled them to lie between 0 and 1. Dashed vertical lines show laser onset and offset. Photocurrent kinetics show slight differences across power during onset, indicated by black arrow. **f**, Simulated example showing the effect of the overapproximation constraint. Left: simulated traces containing a a mixture of photocurrents and PSCs. Right: Photocurrent estimates for the traces shown on the left. The overapproximation constraint allows PhoRC to extract the photocurrent waveform while ignoring PSCs.

Within a given dataset, kinetics were largely conserved across trials (Figure 2a,b with some exceptions, discussed below). This motivated us to consider a low-rank model in which a learned temporal waveform is scaled at each stimulation (Figure 2c). Given *N* recorded traces, each with *T* timesteps, we can express our model as

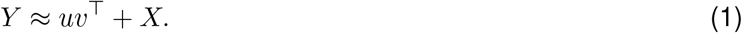

Here, *Y* is an *N* × *T* matrix with recorded traces along its rows, *X* is a matrix capturing evoked PSCs at each trial, and the rank-one term *uv*^⊤^ contains our model of the photocurrent artifact, where *u* is a *N*-length vector containing the scalings, and *v* is a *T*-length vector containing the photocurrent waveform. Due to the short ISIs used in our experiments, we occasionally observed residual photocurrents from prior trials which impacted photocurrent estimation. To handle this, we introduced additional terms which model these effects (section 4.3.2). The model described by Equation 1 is similar to low-rank methods which have been applied to the removal of electrical artifacts from extracellular array recordings [Mena et al., 2017, O’Shea and Shenoy, 2018]. Unlike prior approaches which have relied on fixed templates [Merel et al., 2016, Printz et al., 2021] we propose to automatically learn the photocurrent waveform *v* and scalings *u* from the data itself.

The key challenge in photocurrent subtraction is to separate the photocurrent component *uv*^⊤^ from the PSC component *X*. To this end we introduce two key innovations. First, we use a short window of time during stimulation, the “photocurrent integration window,” to learn the photocurrent scalings *u*. We found that this improved the model’s capacity to separate photocurrent from PSCs, since most PSCs occur after the integration window.

Second, we introduce an “overapproximation” constraint which requires our photocurrent estimate to lie above the recorded trace (Figure 2d, see section 4.3.2 for more detail). This constraint leverages the fact that, compared to photocurrents, synaptic currents are much more variable. Whereas photocurrents only involve the gating of a single molecule (the opsin), synaptic currents depend on a complex array of presynaptic events – principally, the upstream spike and subsequent vesicle release – each with substantial stochasticity. Since photocurrents and PSCs are summed (Figure 1a), the photocurrent waveform present on each trial is an overapproximation of the observed current (Figure 2d). Thus, the overapproximation constraint serves to separate signal from artifact, making it possible to estimate *u* and *v* from the data despite the present of PSCs. Such constraints have been explored in the machine learning literature for robust fitting of low-rank models [Gillis and Glineur, 2010, Tepper and Sapiro, 2016], and have previously been applied when modeling barcode fluorescence data [Chen et al., 2022]. Our approach is similar to the asymmetric loss function used when processing calcium imaging data in refs. [Inan et al., 2017, Inan et al., 2021].

Two additional steps were necessary in order for the low-rank method to achieve good performance on real data. Up to this point, we have assumed that photocurrent kinetics are constant across all recorded traces from a given experiment. While we found photocurrent kinetics to be much more consistent within a dataset than between datasets, we did observe slight power-dependent changes in kinetics within a given experiment. To understand this effect, we selected representative photocurrent traces from a single ChroME2f experiment (Figure 2e, left traces). We then averaged traces collected at a single laser power, and scaled them to lie between zero and one, yielding a template of photocurrent kinetics at each power (Figure 2e, right traces). We found that, at higher powers, the photocurrent reaches its peak value slightly faster. Though this effect is small, it is important to consider for our purposes – even small errors in photocurrent subtraction can leave residuals which impact connectivity estimation. To account for these changes, we first sorted all recorded traces by the magnitude of the current during the stimulation window. We then ran the low-rank model separately on small batches of sorted traces (typically 100 at a time). By using this batched strategy, we encourage photocurrent kinetics to be conserved within a given batch. With this batching strategy in place, the low-rank model is better able to capture the photocurrent waveform.

Finally, we sometimes found qualitatively that a rank-two model was a better fit for the data than the rank-one model above – i.e., photocurrents are close to but not exactly scaled copies of each other (see Methods 4.3). In practice, it is straightforward to select the rank of the model by subjective evaluation; we never found the need to use more than two components.

We found that the low-rank model was computationally efficient, removing photocurrents from a 25 minute experiment in under four minutes when running on a laptop. This faster than real time performance creates the potential for future online applications (see section 3). We refer to the entire procedure for removing photocurrents as Photocurrent Removal with Constraints (PhoRC).

### 2.3 PhoRC captures photocurrent artifacts across protocols and datasets

We first sought to qualitatively evaluate the estimates from PhoRC and ensure they successfully captured our intuition about which signals should be captured and which should be ignored. Using the same targeted and grid datasets shown in Figure 1, we used PhoRC to subtract photocurrents and subsequently estimated connectivity using the CAVIaR pipeline [Triplett et al., 2022]. As expected, we found that PHoRC resulted in pruned connections (Figure 3a, left panels) and reduced connection strengths (Figure 3a, right panels) for targets with photocurrent contamination. For the large-scale experiments we performed, qualitative evaluation can be difficult as it is impractical to manually inspect the thousands of traces collected during a single experiment. We therefore found it useful to assess performance on two categories of traces: those with the largest putative photocurrent response, and those with the largest putative synaptic current response (see subsection 4.5). In examining these representative traces, we found that PhoRC inferred the photocurrent waveforms while ignoring PSCs for both the grid and targeted protocols (Figure 3b). This was especially evident in the targeted dataset, in which some traces showed a mixture of photocurrents and putative PSCs (Figure 3b, right two panels).

**Figure 3:**
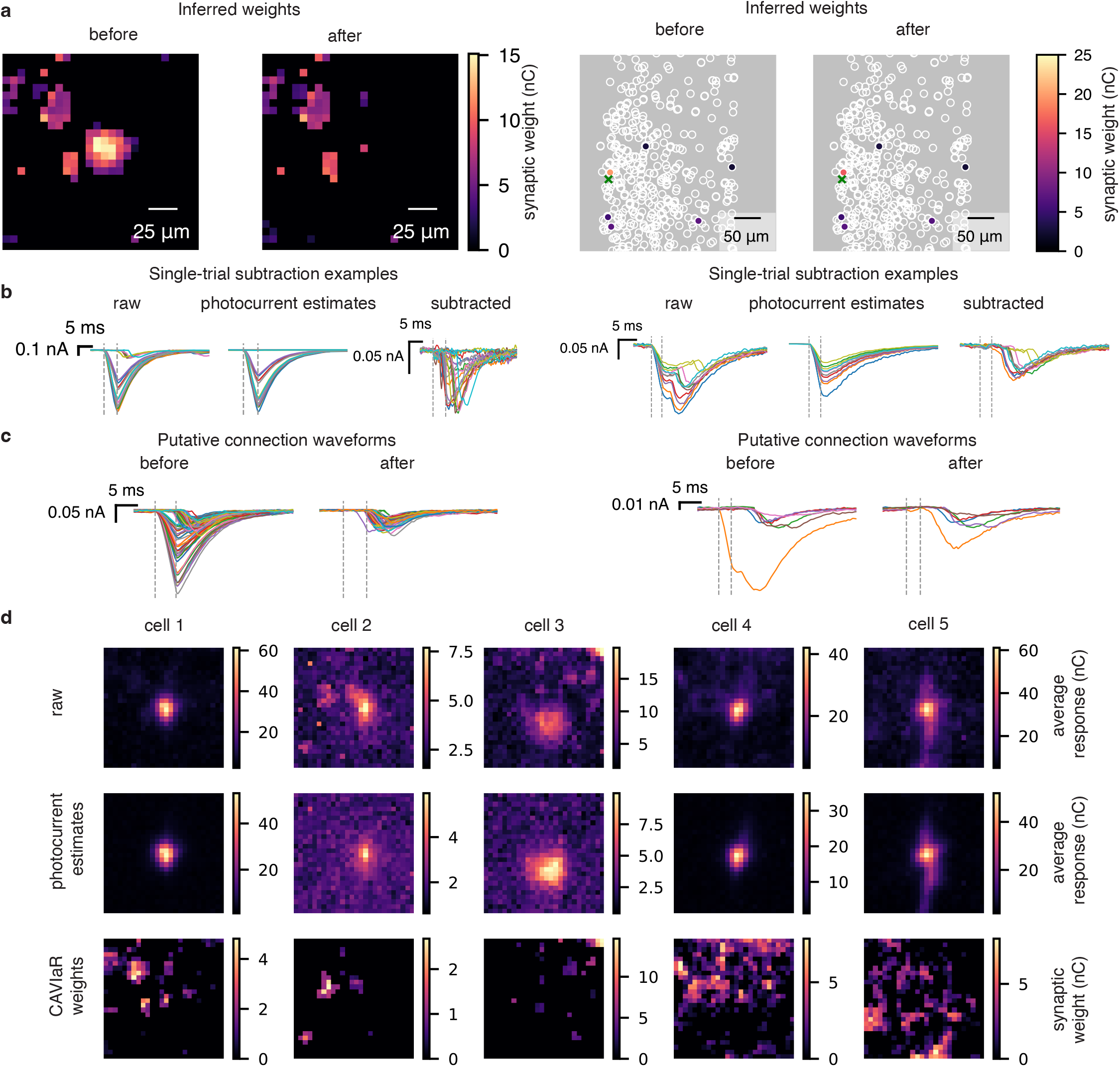
PhoRC captures photocurrent artifacts across protocols and datasets. **a**, Left: Inferred weights for the grid design, before and after subtraction. Weights are shown for a single Z plane. Right: Inferred weights before and after subtraction for a targeted experiment. Weights are shown for the entire Z stack. Green cross marks location of the postsynaptic cell. Applying PhoRC reduced the inferred synaptic strength of the connection nearest the postsynaptic cell, which was contaminated by photocurrent. **b**, Traces selected by taking 10 trials with the largest estimated synaptic current components, and ten trials with the largest estimated photocurrent components. Left panels: Observed currents, photocurrent estimates, and subtracted traces for a ChroME2f experiment. Right panels: same as in left panels but for a ChroME2s experiment. PhoRC successfully infers the photocurrent waveform in both cases, despite the use of different opsins. Estimates ignore PSCs which have higher latency than photocurrents. **c**, Left panels: average evoked waveforms for putative connections before and after subtraction for a ChroME2f experiment. Right panels: same as left, but for the ChroME2s experiment. In both cases, connection waveforms before subtraction display signs of photocurrent contamination, whereas waveforms obtained after subtraction have PSC-like profiles. **d**, PhoRC performance across grid datasets. Each column shows a different dataset. Responses are averaged across planes. From top to bottom: raw maps, photocurrent estimates, and inferred weights from the CAVIaR pipeline.

Following connectivity inference, we averaged all evoked responses at the highest laser power for the targets labeled as connected. This yields a putative connection waveform which describes the average evoked current when a given target is successfully stimulated (see subsection 4.4). With no subtraction, many of these putative connections displayed waveforms typical of photocurrent, whereas after the subtraction step, these waveforms displayed profiles typical of EPSCs (Figure 3c). In some cases, the uncorrected waveforms were indicative of combined EPSCs and photocurrents (Figure 3c, right two panels). In these cases, the subtracted waveforms maintained their putative EPSC components while the photocurrent components were removed. The subtracted waveforms were also smaller in magnitude, reflecting the fact that, in our experiments, evoked photocurrent artifacts were often much larger than typical PSCs (0.3 - 0.8 nA).

For five datasets collected using the grid mapping protocol, we used PhoRC to subtract photocurrents and subsequently inferred connectivity using CAVIaR (Figure 3). Each dataset was dominated by a prominent peak of recorded current near the center of the grid, the putative location of the postsynaptic cell (Figure 3d, top). By averaging the the inferred photocurrent response at each pixel, we obtained spatial maps of evoked photocurrents (Figure 3d, middle). The scale of this photocurrent response varied signficantly between datasets, likely due to variation of opsin expression in the postsynaptic cell. After subtracting the inferred photocurrents, we recovered plausible connectivity maps which were previously obscured by the photocurrent artifact (Figure 3d, bottom). Figures S1-S5 show representative traces and the corresponding subtraction results for each of the five datasets displayed in Figure 3d, and Figure S6 shows additional results for an experiment in which the postsynaptic cell is a putative parvalbumin-expressing (PV) neuron. By examining the recorded current traces and photocurrent estimates, we found that PhoRC was able to adapt to the subtle kinetics differences we observed between datasets.

### 2.4 Validating PhoRC pharmacologically and via simulated data

#### 2.4.1 Validating PhoRC on real datasets using synaptic blockers

A critical difficulty in developing any tool for photocurrent subtraction is the lack of ground-truth data. We therefore sought to further validate PhoRC through pharamcological blockade of synaptic signaling. We first patched an opsin positive cell and mapped incoming connections using the grid protocol (Figure 4a). As expected, recorded traces contained a mix of synaptic currents and photocurrents during this control block (Figure 4b).

**Figure 4:**
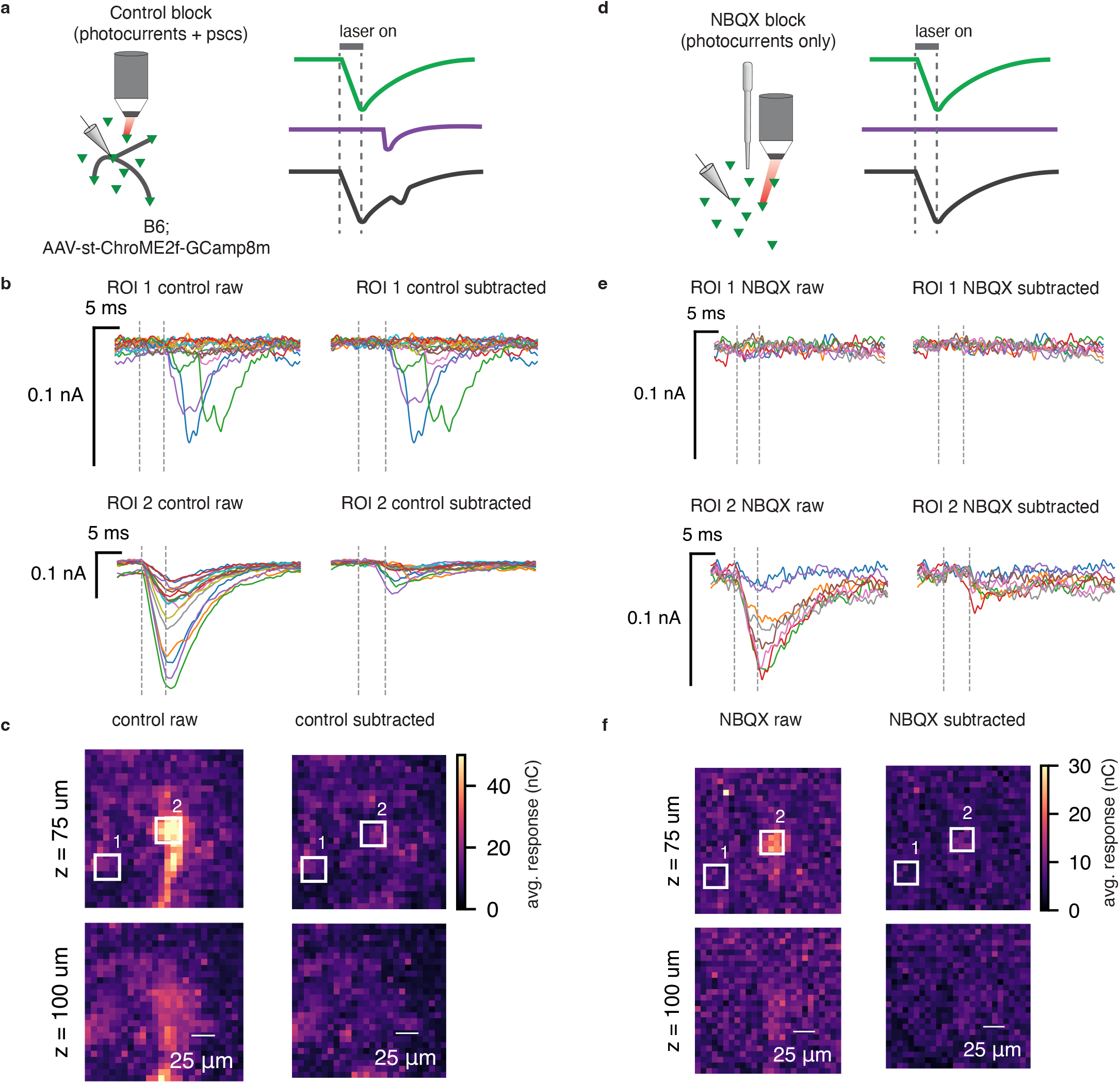
Validating photocurrent subtraction with synaptic transmission block. **a**, We used the grid design to map synaptic connections in a two part experiment. During the control block, we recorded photocurrents along with synaptic currents as in prior experiments. **b**, Raw and subtracted traces from two ROIs during the control block. ROI 1 is far from the putative location of the postsynaptic cell, while ROI 2 is nearby. **c**, Raw and subtracted grid maps from the control block. The colorbar for the “control raw” map has been truncated to improve legibility. ROIs correspond to the traces shown in **b. d**, After bath application of NBQX, we repeated the mapping experiment on the same patched cell, recording isolated photocurrents. **e**, Raw and subtracted traces from the NBQX block for the same two ROIs as in **b**,**c**. As expected, nearly all currents are removed from ROI 2. **f**, Raw and subtracted grid maps from the NBQX block. As expected, the subtracted maps during the NBQX block appear blank. ROIs correspond to the traces shown in **e**.

Following the control block, we bath applied the AMPA receptor antagonist NBQX to block synaptic transmission (NBQX block, Figure 4d), and observed that the EPSCs visible during the control block were removed. If our photocurrent estimates are accurate, we would expect that PhoRC should remove all currents present during the NBQX block. Indeed, we found that nearly all currents were subtracted, resulting in a blank map devoid of photocurrents and synaptic currents (Figure 4e,f).

#### 2.4.2 Validating PhoRC using simulated mapping experiments

We next sought to understand how PhoRC affects connectivity inference. To do so, we used simulated datasets in which ground truth responses and connectivity were known. We simulated mapping experiments using a population of 100 upstream targets with ten percent connection probability. A fraction of targets, chosen at random, evoked direct photocurrent when stimulated (Figure 5a). Photocurrent waveforms were simulated using the model from ref. [Schoeters et al., 2021], with parameters chosen at random to capture a wide variety of photocurrent kinetics (Figure 5e vs f, raw traces). Our simulated mapping experiments include a great deal of biological detail not directly present in the photocurrent removal model, including power-dependent PSC latency and spontaneous PSCs.

**Figure 5:**
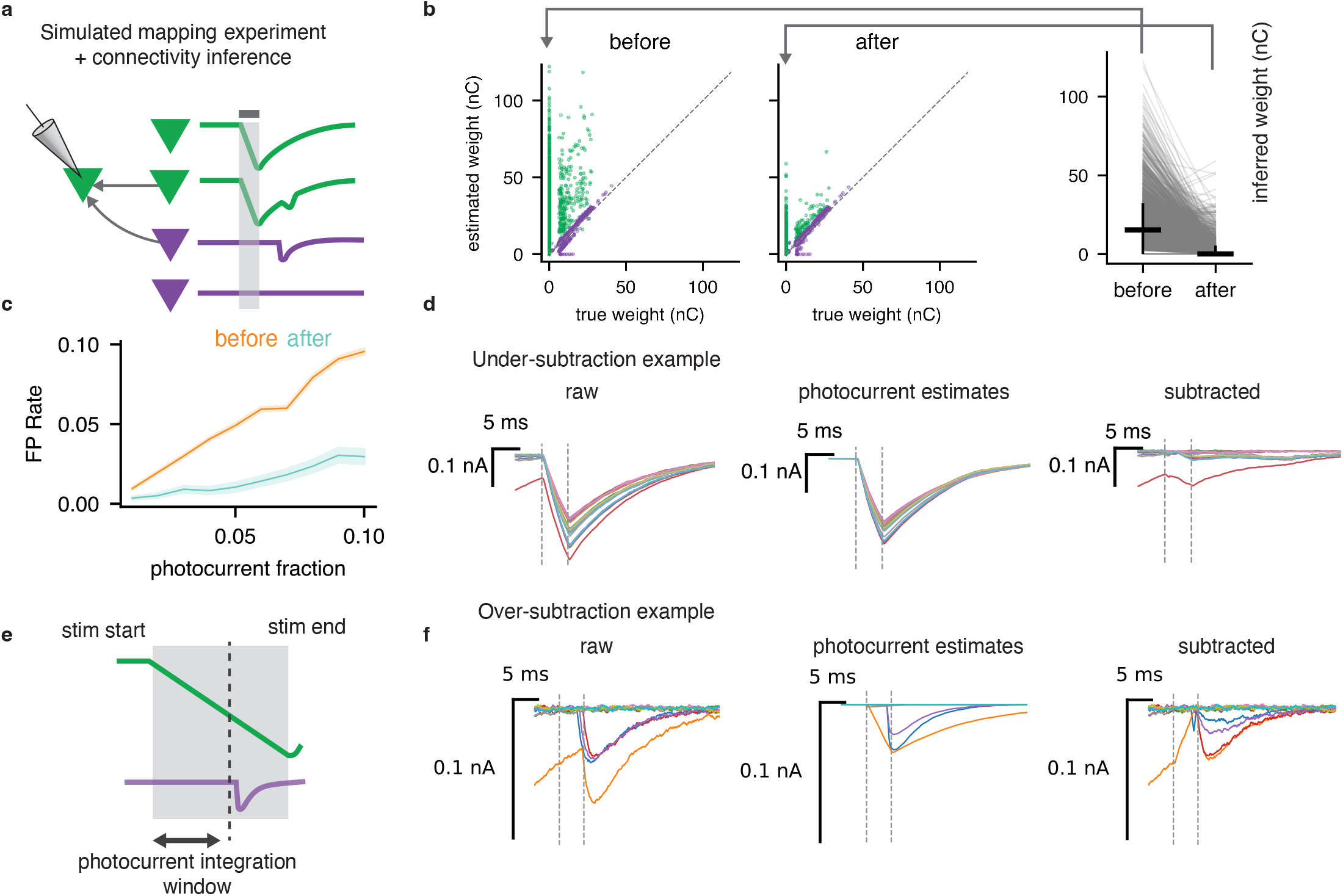
Validating photocurrent subtraction with simulated mapping experiments. **a**, We simulated complete circuit mapping experiments in which stimulating some cells (green) evoked direct photocurrent. We then used a connectivity inference pipeline to assess how photocurrent subtraction affected connectivity inference. **b**, Left two panels: scatterplots of inferred vs. true weights before and after photocurrent subtraction. Vertical axis shows estimated weight when connectivity is inferred after subtraction, horizontal axis shows estimated weights with no photocurrent. After subtraction, the photocurrent-inducing cells (green) are pushed closer to the identity line. Right panel: before vs. after for cells which are unconnected yet evoke photocurrent (left-most green dots in b). Vertical bars = standard deviation, horizontal bars = median. **c**, False positive rate as a function of the number of cells which cause direct photocurrents before (orange) and after (blue) subtraction. **d**, Traces from an example false positive in which residual current after subtraction caused us to falsely infer a connection. This typically occured due to residual photocurrent from prior trials. **e**, The photocurrent integration window is used by PHoRC to estimate the amount of photocurrent present in each trace. This parameter allows the user to trade off between more aggressive subtraction (larger integration window) at the expense of occasional over-subtraction, as shown in **f. f**, Example of over-subtraction that can occur when the integration window extends beyond the minimum PSC latency.

We swept the number of photocurrent inducing cells (“photocurrent fraction”) from one to ten percent and subsequently inferred connectivity. For each value of the photocurrent fraction, we used 10 random realizations of a simulated mapping experiment, in which each realization had random assignments of connectivity and photocurrent kinetics (see subsection 4.6). We found significant improvement in connectivity estimates after applying PhoRC (Figure 5b,c). Without applying any subtraction method, the number of false positive connections scaled linearly with the photocurrent fraction (Figure 5c). Applying PhoRC to estimate and subtract photocurrents resulted in signficantly fewer false positive connections while preserving nearly all true positive connections (Figure 5b,c).

For unconnected cells which induced direct photocurrent, the median inferred synaptic weight after applying PhoRC was zero (Figure 5b, rightmost panel). Nonetheless, the subtraction step is not perfect. We did observe a small number of false positives among unconnected photocurrent-inducing cells (Figure 5b,c). In most cases, these false positives were caused by residual effects from the prior trial (example shown in Figure 5d). Although our model contains terms which account for this effect, these cases still present a challenge.

Our simulations also demonstrated that in rare cases, PhoRC can partially subtract PSCs. The key parameter governing this behavior is the integration window. As detailed in section 4.3.2, PhoRC proceeds in two steps. In the first step, PhoRC uses a small window of data (typically the period while the laser is on) to learn the amount of photocurrent present in each trace. We term this slice of the data the integration window, shown graphically in Figure 5e. Our simulations showed that this parameter controls a tradeoff between more aggressive subtraction (longer window) and more conservative subtraction (shorter window). We noticed that when the integration window extends beyond the minimum PSC latency, PhoRC will occasionally “over-subtract” and partially remove low-latency PSCs. For the simulation results in Figure 5b-d, we therefore set the integration window to be three milliseconds, matching a typical value for low-latency PSCs. We expect that this value for the integration window will work well in most cases, with longer windows only necessary in cases of extreme photocurrent contamination.

### 2.4 PhoRC preserves real PSCs in hybrid simulated-plus-real datasets

To further validate PhoRC, and assess the impact of varying the integration window, we used “hybrid” datasets in which simulated photocurrents were added directly to recorded EPSCs. Such an approach to validation has become common practice in the spike-sorting literature, where ground-truth datasets are similarly difficult to obtain [Rossant et al., 2016, Pachitariu et al., 2016]. The hybrid dataset approach presents a more realistic challenge than purely synthetic data for two reasons. First, the distribution of PSC latencies more closely matches what we expect in real experiments. Second, the ground-truth recordings contain electrical noise which may not have been fully captured by our simulations.

For this set of experiments, we used the sparsely-expressing Ai203 line [Bounds et al., 2021], and performed grid mapping while recording from opsin-negative cells. We then added synthetic photocurrents to these datasets, and tested our ability to recover ground-truth EPSCs (Figure 6a).

**Figure 6:**
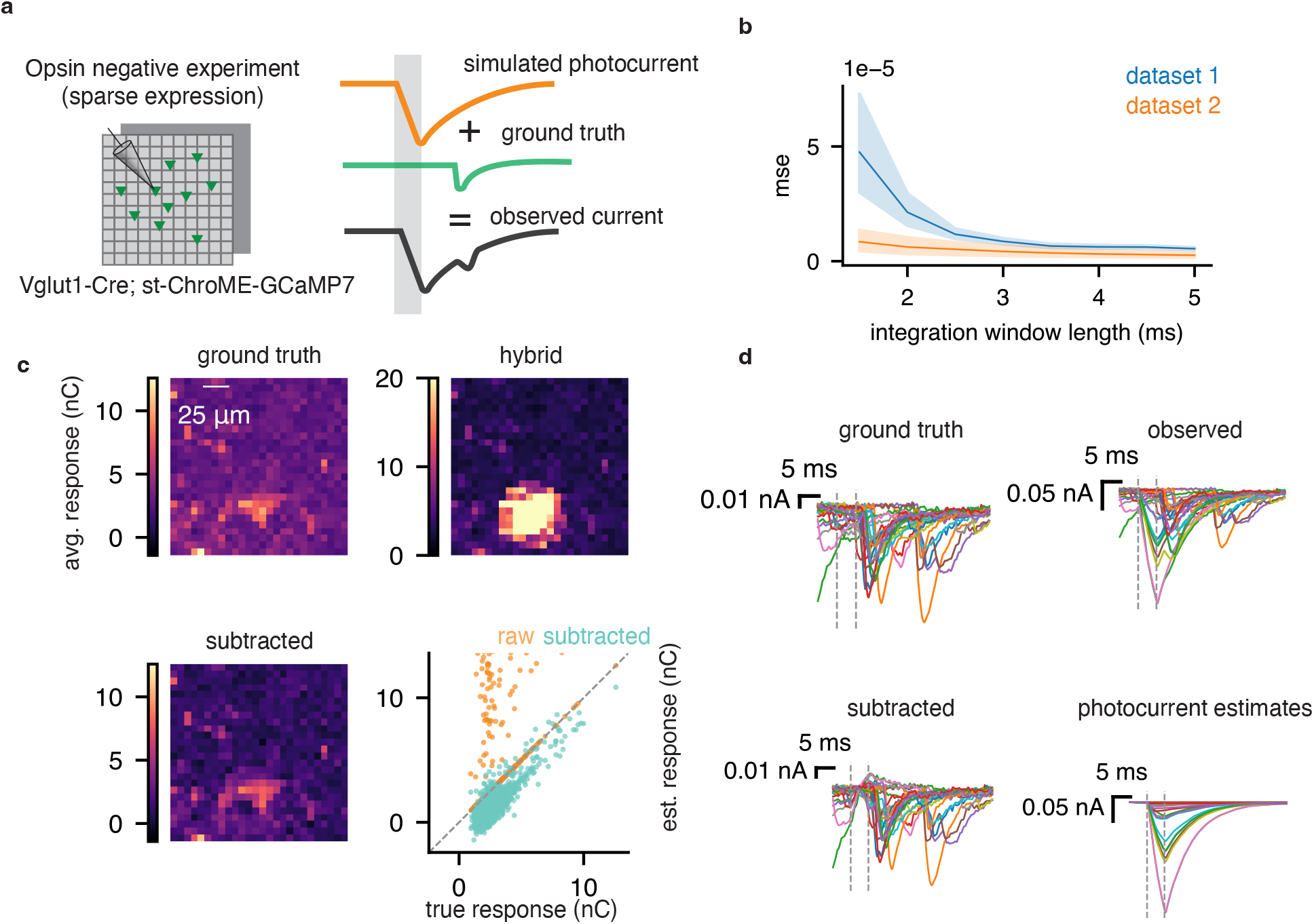
Validating photocurrent subtraction via simulated-plus-real “hybrid” datasets. **a**, Using a sparsely expressing preparation, we mapped synaptic connectivity using the grid design. We then added simulated photocurrents to these recorded EPSCs in order to validate the low-rank model. **b** Mean-square error (MSE) between ground-truth PSCs and estimated PSCs for varying values of the integration window. We see diminishing returns when using windows longer than 3 milliseconds. **c** Ground truth EPSC map, observed map containing real EPSCs and simulated photocurrents, and map after subtraction. Bottom right shows the true vs. estimated responses, before and after applying PhoRC. **d**, Example traces corresponding to the maps shown in **c**. Subtracted EPSCs closely mirror ground-truth.

An example hybrid dataset is shown in Figure 6c,d. We simulated photocurrent contamination by adding spatially-varying photocurrent responses, in which the mean evoked photocurrent decayed with distance from the patched cell. In order to make the experiment more challenging, we deliberately placed the peak of photocurrent contamination on top of an existing connection. After applying PhoRC, we recovered the spatial structure of the ground-truth connectivity map (Figure 6c), and our recovered synaptic responses closely matched the ground-truth responses (Figure 6c,d).

Since our simulations demonstrated the importance of the integration window parameter, we used the hybrid dataset approach to measure this parameter’s impact on performance. For two opsin-negative datasets, we added synthetic photocurrents and measured the mean-square error (MSE) between the true and recovered synaptic currents. As we expected, the MSE decays with increasing window lengths, because more photocurrents are captured (Figure 6b). The MSE reaches its minimum value for window lengths around 3 milliseconds, suggesting that this value may be a good starting point for practitioners, and we used this value for the example experiment shown in Figure 6c,d.

### 2.5 PhoRC is compatible with accelerated compressed sensing experiments

Until this point, we have considered mapping experiments in which a single upstream target is probed at a time [Baker et al., 2016, Izquierdo-Serra et al., 2018, Hage et al., 2022]. However, a growing body of evidence suggests that mapping can proceed faster if a random ensemble of upstream cells is stimulated simultaneously, and connectivity is reconstructed using computational demixing approaches such as compressed sensing [Hu et al., 2009, Packer et al., 2012, Shababo et al., 2013, Draelos et al., 2020, Triplett et al., 2022]. We refer to this experimental setup as ensemble stimulation.

In the case of ensemble stimulation, photocurrent contamination has even larger effects on synaptic mapping – if even one stimulated target per ensemble induces direct photocurrent, the entire measurement will be corrupted (Figure 7a,e). Thus even cells which are far from the patched cell may have corrupted connectivity estimates. We therefore asked whether PhoRC can still remove photocurrents in the case of compressed sensing experiments.

**Figure 7:**
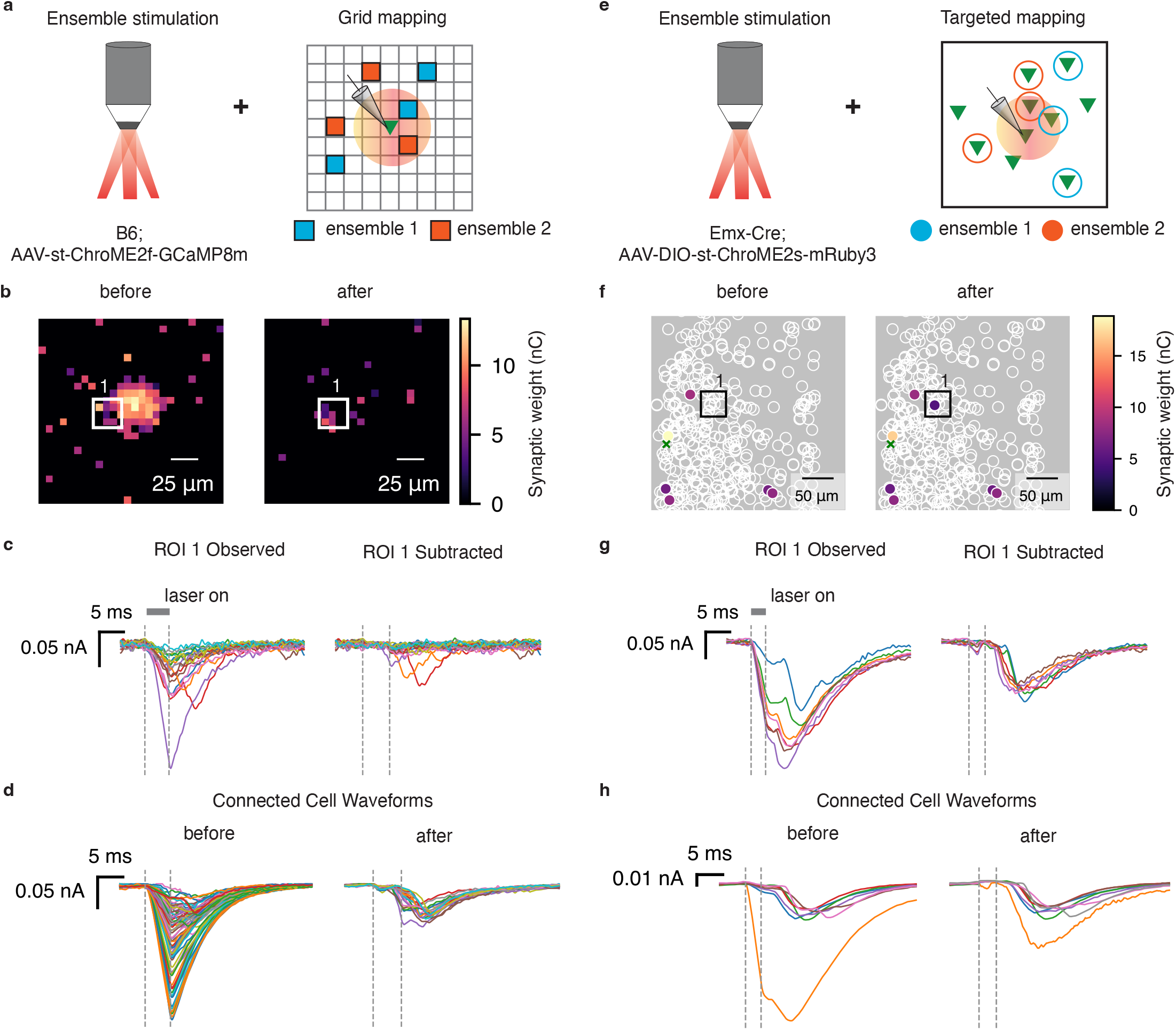
PhoRC allows ensemble mapping in the presence of photocurrent artifacts. **a**, Cartoon depicting the photocurrent contamination problem with ensemble grid stimulation. Red circle denotes region where stimulation will evoke direct photocurrent. If one grid point in an ensemble evokes photocurrent, the entire measurement will be corrupted. **b**, Inferred grid connections before and after photocurrent subtraction. ROI denotes location of a putative connection. **c**, Observed and subtracted traces from the ROI selected in **b**. Traces from this ROI contain direct photocurrents whenever a different target in the ensemble evokes direct photocurrent. **d**, Inferred waveforms for each putative connection before and after subtraction. **e-h**, Same as **a-d** but for the targeted mapping design. **e**, Cartoon depicting the photocurrent contamination problem with ensemble targeted stimulation. As in **a**, red circle denotes region where stimulation evokes direct photocurrent. **f**, Schematic of inferred connectivity before and after subtracting photocurrents, Z projection. ROI corresponds to a region far from the postsynaptic cell, but photocurrents are still present due to the use of ensemble mapping. **g**, Observed and subtracted currents from the ROI shown in **f. h**, Inferred waveforms for each putative connection before and after subtraction.

For the postsynaptic cells shown in Figure 1 and Figure 3, we additionally performed accelerated mapping with both the grid and targeted protocols in which ten targets were simultaneously stimulated through holography. Qualitatively, photocurrents induced by 10-target mapping closely resemble those induced during single-target mapping (Figure 7c,g, observed traces). As in the single-target mapping case, we observed relatively consistent photocurrent kinetics within a given dataset, allowing PhoRC to effectively extract the photocurrent waveform. For the grid dataset shown in Figure 7, we additionally show a comparison of synaptic weights obtained via single-target and ensemble mapping in Figure S1.

We used the pipeline from ref. [Triplett et al., 2022] to reconstruct connectivity both before and after applying PhoRC to subtract photocurrents. As before, photocurrent subtraction pruned many inferred connections near the patched cell (Figure 7b, left panels). However, in this case, we also observed that PhoRC removed photocurrents on trials which included targets far from the postsynaptic cell (Figure 7f,g). Due to the use of compressed sensing, we occasionally observed that application of PHoRC resulted in the addition of new connections (Figure 7b,f). This reflects the fact that a large fraction of recorded traces contained photocurrent in these experiments, due to the use of ensemble stimulation (Figure 7a,e), and removal of these currents can alter the behavior of compressed sensing algorithms. As in prior experiments, we found that PhoRC effectively suppressed photocurrents while preserving PSCs, and adapted accurately to the different photocurrent kinetics of the opsins used in the grid vs. targeted experiments (Figure 7c, g). Before applying PhoRC, many inferred connections displayed average evoked waveforms resembling photocurrents, whereas after applying PhoRC, the average waveforms matched the typical profile of EPSCs (Figure 7d, h).

## 3 Discussion

We developed a matrix-factorization based method called Photocurrent Removal with Constraints (PhoRC), which subtracts photocurrent artifacts while preserving synaptic current signals. In our experiments, we used soma-targeted expression of ChroME2s and ChroME2f, which reduces expression in neurites and thereby limits the region proximal to a cell of interest in which off-target stimulation evokes direct photocurrent [Shemesh et al., 2017, Mardinly et al., 2018, Sridharan et al., 2022]. Nonetheless, we observed confounding photocurrent artifacts when using two-photon holographic optogenetics to map synaptic connections near the postsynaptic cell. After applying PhoRC, we were able to map these connections despite confounding photocurrent effects. PhoRC was able to infer the photocurrent waveform for two opsins with different kinetics, and subsequently improved the accuracy of two approaches for synaptic mapping (grid-based and target-based) and across both simulated and hybrid datasets. Via our synaptic blocker validation experiment, we verified that PhoRC is able to achieve near-complete subtraction of photocurrents.

Finally, we found PhoRC to be compatible with compressed-sensing based mapping experiments in which ensembles of candidate presynaptic cells are excited simultaneously. Compatibility with compressed sensing is crucial given recent evidence that such techniques can accelerate mapping by nearly an order of magnitude [Draelos et al., 2020, Triplett et al., 2022]. PhoRC is especially applicable to compressed sensing experiments given that the probability of observing the photocurrent artifact grows with the number of targets in the ensemble. In our ensemble mapping experiments, we found that PhoRC removed confounding photocurrents even for ensembles which contained targets far from the postsynaptic cell. Thus, we predict that the combination of PhoRC with compressed sensing will allow mapping of both near and far connections onto a single neuron.

As with any computational tool, some care must be taken when interpreting the outputs of PhoRC. In some cases, PhoRC can leave residual photocurrents, leading to false positive connections. Although the model includes terms which account for residual photocurrent from prior trials (see section 4.3.2), we found in simulation that these residual effects could present challenge cases for PhoRC. These cases are more common when using short inter-stimulus intervals and slower opsins.

There has been limited prior work on removing the photocurrent artifact. Printz *et al* manually selected traces which contained only photocurrent, and averaged these traces to extract a separate photocurrent waveform for each presynaptic target [Printz et al., 2021]. The authors also noted that they were forced to discard data in which the artifact was larger than a certain threshold. This manual strategy implicitly assumes that photocurrent amplitude is identical between successive stimulations of the same presynaptic target. By contrast, the low-rank model used here adapts seamlessly to random variation in photocurrent amplitude.

Merel and Shababo *et al* developed a Bayesian framework in which photocurrents could be removed given knowledge of the photocurrent waveform; however, this approach requires *a priori* knowledge of photocurrent kinetics, and assumes that these kinetics are independent of applied laser power [Merel et al., 2016]. Unlike these prior approaches, PhoRC learns the photocurrent waveform from the data itself and can adapt to power-dependent changes in the temporal shape of the photocurrent, allowing practitioners to use their entire dataset without time-consuming manual removal of artifacts.

One other related approach is that of [Triplett et al., 2022], which used a non-linear signal processing technique called neural waveform demixing (NWD). NWD uses a neural network trained on simulated data to denoise recorded PSC traces and correct for confounded baselines. In some regimes, NWD can effectively separate PSCs from photocurrents due to the difference in latency. In our experiments, we found that NWD reduced the impact of photocurrents, but did not eliminate them. Indeed, all connectivity estimates shown above (e.g Figure 1d,g) used NWD as a preprocessing step before connectivity inference.

In the process of developing PhoRC, we also explored the possibility of using a neural network (a la NWD) to pool information across recorded traces and thereby extract the photocurrent waveform [Zaheer et al., 2017]. While this approach worked well, we found that it required costly re-training of networks for different stimulus durations. We therefore adopted the low-rank model described here, which achieved similar accuracy and adapts easily to different pulse durations without requiring any laborious neural network retraining. In certain cases, the noise-robustness of a neural network approach may be preferred, and we leave this as an avenue for future work.

Several possible extensions of PhoRC may increase its applicability. First, while we demonstrated post-hoc removal of photocurrents following the completion of an experiment, it may be useful in future applications to remove photocurrents while an experiment is in-progress. Running on a laptop, PhoRC runs faster than real time and is therefore well suited to online applications. This creates the potential for adaptive experiments in which practitioners are able to immediately identify putative connections amidst photocurrent artifacts, and target these putative connections for further study. Finally, while we have considered mapping experiments in acute slices, *in vivo* connectivity mapping might provide deeper insights into the circuit mechanisms engaged during behavior. The *in vivo* setting may require extensions of PhoRC to handle increased spontaneous activity and more frequent recording instability.

We view PhoRC as one method in an increasingly broad suite of computational tools which allow two-photon optogenetics to thoroughly characterize monosynaptic connectivity [Hu et al., 2009, Draelos et al., 2020, Triplett et al., 2022, Triplett et al., 2023]. PhoRC is compatible with methods for accelerated mapping via compressed sensing, and thus we predict it will enable experimenters to measure both short and long-range connectivity in a single experiment. Such experiments would contribute to a more complete view of neural connectivity, and could subsequently inform improved models of neural circuits.

## Acknowledgements

We thank Darcy Peterka for helpful comments and discussions. This work was funded by NIH awards 1RF1MH120680 and 1U19NS107613-01 to HA and LP. BA, MAT, and LP were supported by the Gatsby Charitable Foundation and NSF NeuroNex award 1707398.

## 4. Methods

### 4.1. Experimental Methods

#### 4.1.1 Animals

The mice used for experiments in this study were C57BL/6 (B6; JAX stock#000664) and Emx1-Cre (JAX stock# 005628), with the latter further crossed with CD-1 (ICR; Charles River) mice. Mice were housed in cohorts of five or fewer in a reverse light:dark cycle of 12:12 hours, with experiments occurring during the dark phase. All surgical procedures and experiments on animals were conducted in compliance with the Animal Care and Use Committee of the University of California, Berkeley. All experiments used male and female mice equally.

#### 4.1.2 Viral expression of opsins

B6 or Emx-Cre mice were neonatally injected with adeno-associated virus (AAV)-driven vectors to pan-neuronally express Chrome2f via the synapsin promoter (AAV9.hSYN.ChroME2f-6xHIS.GCaMP8m-ST.SV40) or selectively express Chrome2s in excitatory neurons in a cre-dependent manner (AAV9.CAG.DIO.ChroME-ST.P2A.H2B-mRuby6)), respectively. Both custom made viral preparations for Chrome2f and 2s were generated by the Penn Vector Core. Pups aged P3-5 were anesthetized by placing them on ice for approximately 3 minutes. Each animal was then stabilized and virus was injected using a Nanoject III nanoliter injector (Drummond Scientific Company) in 4 sites surrounding the area of V1. For each site, virus was injected with 30 nl per injection at 5 depths approximately spanning the cortical layers. After injections, the animal was placed on a heating pad for post-surgical recovery. All injected pups were returned to home cage with parent mouse and housed together until weaning age (P21).

#### 4.1.3 3D-SHOT holography

All in vitro electrophysiology experiments employed 3D scanless holographic optogenetics with temporal focusing (3D-SHOT), as described previously [Mardinly et al., 2018, Sridharan et al., 2022, Triplett et al., 2022]. Based on the Movable Objective Microscope (MOM; Sutter Instrument Co.) platform, the setup is built with three combined optical paths: a 3D two-photon (2p) photostimulation path, a fast resonant-galvo raster scanning 2p imaging path, and a widefield one-photon (1p) epifluorescence/IR-transmitted imaging path, merged by a polarizing beamsplitter (PBS) before the microscope tube lens and objective. A Monaco 1035-80-60 737 (1040nm, 1MHz, 300fs, Coherent Inc.) fiber laser was used for photostimulation and a Mai Tai Ti:sapphire laser (Spectra Physics Inc.) was used for 2p calcium imaging. Temporal focusing of the photostimulation beam from the femtosecond fiber laser was achieved with a blazed holographic diffraction grating (33010FL01-520R Newport Corporation). The beam was then relayed through a rotating diffuser to randomize the phase pattern and to expand the temporally focused beam to cover the area of the high-refresh-rate spatial light modulator (SLM; HSP1920-1064-HSP8-HB, 1920 × 1152 pixels, Meadowlark Optics). Holographic phase masks were calculated using the Gerchberg-Saxton algorithm and displayed on the SLM to generate multiple temporally-focused spots in 2D or 3D positions of interest [Mardinly et al., 2018]. The photostimulation path was then relayed into the imaging path with a PBS placed immediately prior to the tube lens. As described in [Mardinly et al., 2018], to limit imaging artifacts introduced by the photostimulation laser, the photostimulation laser was synchronized to the scan phase of the resonance galvos using an Arduino Mega (Arduino), gated to be only on the edges of every line scan.

### 4.2 In vitro whole-cell electrophysiology

In vitro slice recordings were performed on 300*µ*m-thick coronal slices obtained from both male and female mice aged P24-40. Slice preparation followed previously described methods [Pégard et al., 2017, Bounds et al., 2021, Triplett et al., 2022]. For each mouse brain to be prepped, opsin expression was checked using a handheld laser light to visualize fluorescence of the mRuby reporter (red; for Chrome2s) or fused GCaMP8 (green; for Chrome2f). Whole-cell patch-clamp protocols were performed in with heating-controlled (33°C) standard ACSF bath solution (in mM: NaCL 119, NaHCO3 26, Glucose 20, KCl 2.5, CaCl 2.5, MgSO4 1.3, NaH2PO4 1.3). Patch pipettes (4-7 MOhm) were pulled from borosilicate glass filaments (Sutter Instrument Co.) and filled with Cesium (Cs2+)-based internal solution (in mM: CeMeSO4 135, NaCl 3, HEPES 10, EGTA 0.3, Mg-ATP 4, Na-GTP 0.3, Qx-314 1, TEA-Cl 5, 295 mOsm, pH=7.45) also containing 50 *µ*M Alexa Fluor hydrazide 488 or 594 dye (ThermoFisher Scientific). Data was recorded at 20 kHz using 700b Multiclamp Axon Amplifier (Molecular Devices). The headstage and electrode holder (G23 Instruments) was controlled by Motorized Micromanipulator (MP285A; Sutter Instrument Co.). All data was acquired and analyzed with custom code written in Matlab using the National Instruments Data Acquisition (DAQ) Toolbox (NI). Membrane (Rm) and series (Rs) resistance were monitored through short sessions of hyperpolarizing steps before and throughout experiments to ensure quality of acquired data. For all experiments, trials with Rs exceeding 30 MOhm were excluded from data analysis. At each stimulation, the response was taken to be the total charge transfer during a 35ms window following stimulation.

#### 4.2.1 Whole-cell grid-based synaptic connectivity mapping

Cells were patched in whole-cell configuration and voltage clamped at -70 mV. Preliminary assessment of photostimulation-evoked postsynaptic responses used widefield 1P stimulation at saturating power. To map synaptic inputs to the patched cell, the surrounding volume of 162.5 × 162.5 *µ*m in lateral (x, y) and 100 *µ*m in axial (z) dimensions was probed through holographic photostimulation of a precomputed three-dimensional grid of holographic spots. Each point (voxel) of photostimulation was spaced 6.5 × 6.5 × 25 (x, y, z) *µ*m apart, targeted randomly at 30 or 40 Hz with 5 ms laser pulses across 20 trials per laser power tested. To build the grid maps, responses at each trial were calculated as the integral of the recorded current for each trial, and theses responses were averaged for all stimulations at a given voxel.

#### 4.2.2. Whole-cell targeted synaptic connectivity mapping

Whole-cell targeted mapping experiments were performed as previously described [Triplett et al., 2022]. An opsin-negative L2/3 putative pyramidal neuron (based on cell body shape) was sealed onto with a recording pipette and the area surrounding the target cell was imaged at 40 frames at 4-5 planes spaced by 25 *µ*m. The positions of the opsin-positive presynaptic candidates were automatically identified in a specified FOV using an in-house algorithm detecting mRuby3 fluorescence in round shapes representing cell nuclei. Next, 10-20 different holograms of either single or ensemble sets of candidate presynaptic targets were computed (see Methods: 3D-SHOT Holography). During hologram computation, stable whole-cell configuration was established in the patched cell and clamped at -70mV. Presynaptic candidates were mapped individually or in 10-target ensembles and across 3-4 powers and repeated 7-15 times per condition in randomly interleaved experimental trials. A map containing 100 presynaptic candidates would contain data from 45 trials of 100 single-target holograms across 3 powers (15 trials/power condition) and 45 trials of 100 different 10-target holograms (100 presynaptic candidates split into 10 different 10-target holograms that were then arranged randomly in 10 sets) across 3 powers (15 trials/power condition), resulting in 495 total trials (45 single-target sets and 450 10-target sets).

### 4.3. The PHoRC algorithm

#### 4.3.1 Unconstrained model via SVD

To build intuition, we begin with a simplified version of the model, and then add constraints and modifications as needed. For simplicity in this section, we reverse the sign of excitatory currents, so that an upwards deflection represents inward current. To begin, assume we have *N* intracellular traces, each with *T* timesteps. We begin by thresholding these traces at zero, since any deflections below zero are due to noise. Let *Y* denote this thresholded matrix *N* × *T*, in which recorded traces are stacked along its rows.

If we plot the rows of the matrix *Y*, we see that the shape of the photocurrent waveform remains largely consistent across stimulations within a single experiment (Figure 2a,b,c). The amplitude of the artifact, however, varies significantly across stimulations, since variable amounts of opsin are excited depending on stimulus location and power. This suggests approximating *Y* using a rank-one matrix, in which a single temporal waveform *v* is scaled at each trial by a parameter *u*_*i*_. We can write this simple model as *Y* = *uv*^⊤^ + *X*, where *Y* is the matrix of observed traces, and *X* is an (unobserved) matrix containing PSCs. Our strategy will be to estimate *u* and *v* in order to ultimately recover *X*. Both *u* and *v* can be estimated by solving the following optimization problem, whose solution is obtained via the SVD of *Y* :

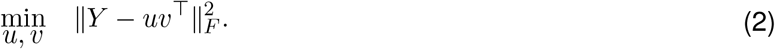

In our experiments, we sometimes observed that photocurrents were not exactly scaled copies of each other. In these cases, we found it beneficial to approximate the photocurrent artifact as a sum of *R* temporal waveforms:

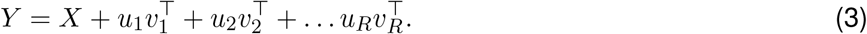

Setting *R* to larger values leads to more complete photocurrent subtraction, at the cost of possible subtraction of PSCs (see section 4.3.6). This model can be compactly written as *Y* = *UV* + *X* where *U* is an *N* × *R* matrix *U*, and an *V* is an *R* × *T* matrix. However, this simple approach fails because the low-rank term *UV* frequently includes PSCs in addition to photocurrents. We therefore introduced constraints which incorporate our prior knowledge about the interaction between PSCs and photocurrents.

#### 4.3.2 Constrained stepwise model

Recall that in this section, we are working with the negative of the recorded traces, so that photocurrents and PSCs are represented as positive deflections from baseline. Since excitatory synaptic currents and photocurrents sum linearly (assuming a perfect space-clamp), the photocurrent waveform present on each trial should be strictly smaller than than the observed trace. We can incorporate this prior knowledge as an under-approximation constraint, which will stop the waveforms *V* from including PSCs. Further, we wish for the weights *U* to encode the strength of the photocurrent artifact at stimulation. If we were to learn *U* using the entire data matrix *Y*, then *U* would include information about the strengths of both photocurrents and PSCs. To avoid this, we instead optimize *U* on a short window of data around laser onset, before most upstream spikes have had time to propagate. These two modifications to the model in Equation 2 give rise to what we term a constrained stepwise model.

We proceed in two stages. First, we fit *U* on a short window of data, which we call the photocurrent integration window. Let *t*_1_ denote the beginning of this window, and *t*_2_ denote the end (see below for how these values can be selected). We learn trial-weights *U*_stim_ by solving

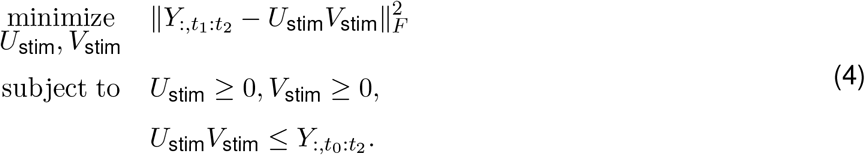

This is a nonnegative matrix underapproximation (NMU) problem. Though it is non-convex, good solutions can be reached efficiently using the alternating directions method of multipliers (ADMM, detailed below). For computational efficiency, we follow the approach used in [Gillis and Glineur, 2010, Tepper and Sapiro, 2016], in which a single rank-one component 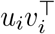 is learned at a time, and then subtracted from the data matrix *Y*. Recursively, we then define 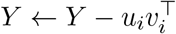and proceed to fit the next rank-one component.

After solving Equation 4, we have obtained values for the photocurrents weight *U*_stim_. We also obtained estimates of the photocurrent waveform(s) *V*_stim_. However, these waveform estimates only account for the short section of data during the photocurrent integration window, from *t*_1_ to *t*_2_. In step two of the PhoRC algorithm, we hold our learned weights constant, and infer waveforms for the entire trial:

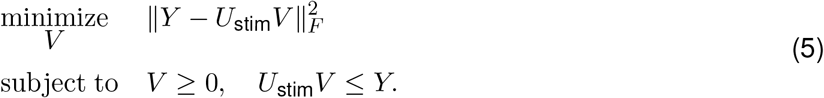

Our estimate of the photocurrent is then *U*_stim_*V*, which we subtract away to yield the underlying EPSCs. Conceptually, these two steps form the core of the PhoRC algorithm. However, we found it helpful to introduce additional constraints in some cases, which we detail below.

### 4.3 3Accounting for prior trial effects and corrupted baselines

Prior work on optogenetic circuit mapping has used relatively long inter-stimulus intervals (ISIs), typically around 100 milliseconds, in order to allow conductances in the postsynaptic cell to return to baseline [Packer et al., 2012, Baker et al., 2016, Hage et al., 2022]. However, [Triplett et al., 2022] demonstrated that it is possible to use much shorter ISIs and subsequently demix the overlapping signals using a neural network.

Up to this point, our model has implicitly assumed that the postsynaptic cell has returned to its baseline at stimulation onset. We found that at the short ISIs used in our experiments (30-33 ms), effects from the prior or subsequent trial could confound photocurrent estimates. This was more common when using ChroME2S due to its slower decay time compared to ChroME2f. To account for this effect, we add a second low-rank component which captures these prior trial effects. Further, we found that some recorded traces had nonzero baselines, which we accounted for using a constant baseline term. With these additional terms, our model can be written as

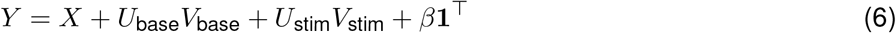

Here, *U*_base_*V*_base_ is a low-rank term which will approximate decaying photocurrents left over from the prior trial. As before, *U*_stim_*V*_stim_ captures photocurrents from the current trial. Finally, *β***1**^⊤^ forms a matrix with constant rows, allowing a nonzero offset for each recorded trace. Finally, we add constraints so that each term in the model behaves as desired. As before, fitting proceeds in two steps, leading to the full PhoRC algorithm described in Algorithm 1. In the first step, we solve the following minimization:

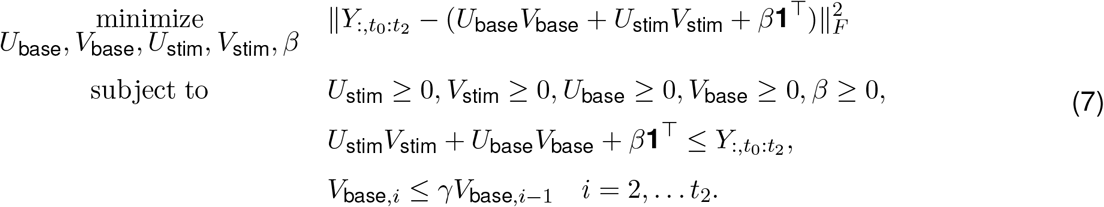

Each constraint has a simple interpretation. The first set of constraints enforces nonnegative weights, waveforms, and offsets. The second set of constraints is the underapproximation constraint, which stops the low-rank terms from including PSCs. In our experiments, we use a rank-one model to capture previous trial effects, so *U*_base_ is a column vector and *V*_base_ is a row-vector.

In the final set of constraints above, *γ* is a hyperparameter between zero and one. Thus, this set of constraints ensures that *V*_base_ decreases exponentially over time, which forces this term to capture baseline effects from the previous trial. The hyperparameter *γ* should be chosen to bound the decay of the opsin, and we found that a value of *γ* = 0.999 worked well for the ChroME opsins used in our experiments. Such a constraint has previously been used to model fluorescence from calcium imaging data, in which an exponential decrease in fluorescence is expected following a spike [Friedrich et al., 2017]. As we will show below, we can therefore leverage optimization strategies originally developed in the context of calcium imaging.

To solve this problem efficiently we use a coordinate descent approach, updating each rank-one term while holding the others fixed. Empirically, we found that only a few (5-10) coordinate descent iterations were needed to obtain good estimates. Full pseudocode for this algorithm is provided in Algorithm 2.

The second step of our algorithm remains conceptually the same as in section 4.3.2. After solving Equation 7, we have obtained values for the relative weights of previous trial photocurrent (*U*_base_), present trial photocurrent (*U*_stim_), and constant baselines (*β*). In step two of the PhoRC algorithm, we hold our learned weights constant, and infer waveforms for both low-rank terms:

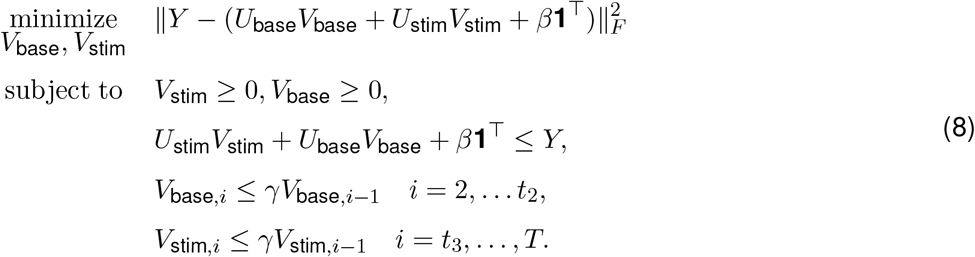

Note that Equation 8 is nearly identical to problem Equation 7. The key difference is that in Equation 7 we use only a short section of the data matrix, and optimize over both weights and waveforms. In Equation 8 we hold weights fixed, using the full dataset to optimize only over the waveforms. The final set of constraints in Equation 8 work to prevent the photocurrent waveform from picking up effects from the *subsequent* trial. The time index *t*_3_ is a user defined hyperparamter that sets the index at which we enforce the exponential decay constraint. In practice, setting *t*_3_ to be 10 milliseconds after stim offset works well. The precise value of this parameter does not have a large impact on results. A schematic showing the role of the parameters *t*_1_, *t*_2_, *t*_3_ is given in Figure S7.

#### 4.3.4 Solving the NMU subproblems

The PhoRC algorithm requires solving several NMU subproblems. In the case where we are not concerned with overlapping trial effects, these problems have the following form:

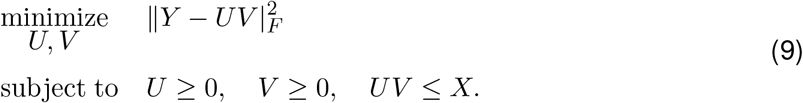

This problem can be solved by fitting a single rank-one component at a time using ADMM. We refer the reader to [Tepper and Sapiro, 2016] for details of this approach. In order to account for prior-trial and next-trial effects, we needed to introduce the exponential decay constraints shown in Equation 7 and Equation 8. Updating a rank-one term in that case results in the following sub-problem:

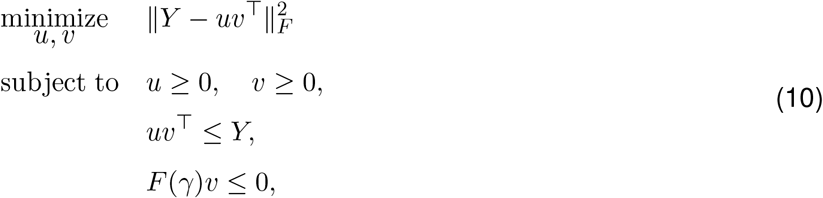

where *F* (*γ*) is a first difference matrix with hyperparameter *γ*:

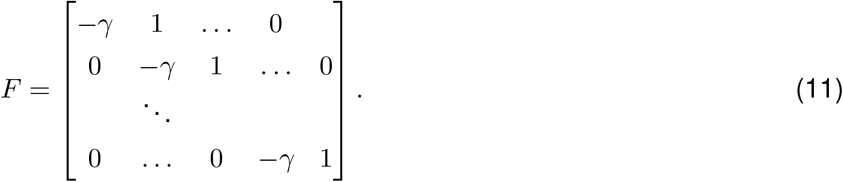

In the case that *γ* = 1, the last constraint in problem 10 becomes a monotonic decay constraint. Our approach will be to optimize simultaneously over *u, v* using ADMM [Boyd et al., 2011]. We begin by rewriting problem 10 using auxiliary variables:

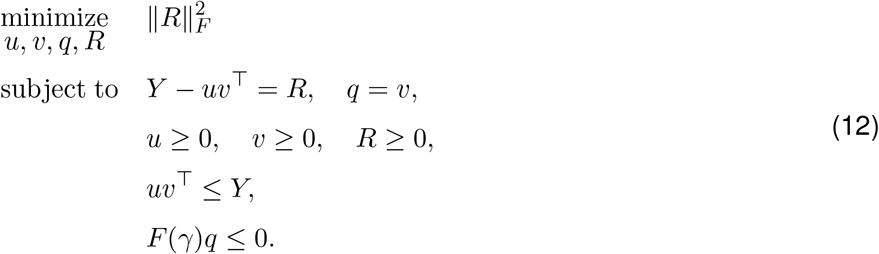

We also introduce dual variables Λ which will be used to enforce the constraint that *Y* − *uv*^⊤^ − *R* = 0 and *λ*, which will be used to enforce the constraint that *q* − *v* = 0. The ADMM udpate for *q* requires the following minimization:

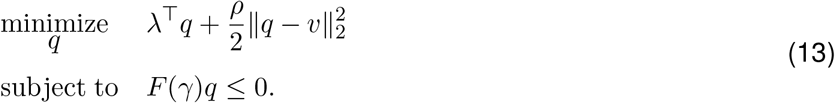

Noting that

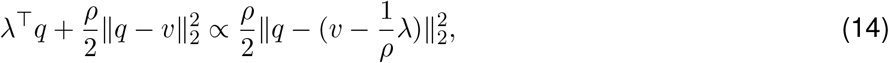

where the proportionality hides terms which are constant with respect to *q*, we can write the above problem as

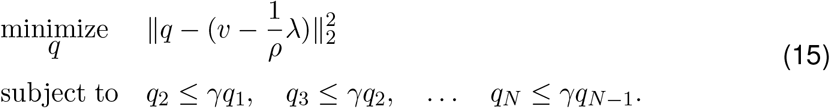

When *γ* = 1 this problem can be solved in 𝒪(*T*) time by the the Pool Adjacent Violators Algorithm (PAVA) [Ayer et al., 1955]. In the case of 0 *< γ <* 1, Friedrich *et al* developed an extension of the PAVA algorithm which efficiently solves this case as well [Friedrich et al., 2017]. The update for *q* has an appealing interpretation: it is a projection of the point 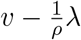 onto the convex set defined by the constraints [Boyd et al., 2011]. When modeling the current-trial photocurrent waveform (*V*_stim_) we are only concerned with enforcing the decay constraint on a subset of the entries. In this case, we simply apply the projection operator to a subset of the entries of *q*.

We can compute the update for *v* similarly. Let *Q* = *Y* − *R*. The update for *v* requires solving the following minimization:

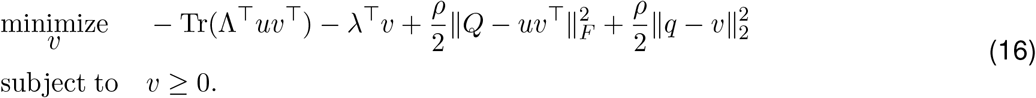

This problem separates along the entries of *v*, and so we can take the gradient of the objective and set it to zero, then threshold the resulting solution. This yields

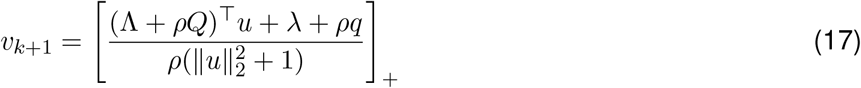

Updates for the other variables are unchanged from the original underapproximation problem considered in [Tepper and Sapiro, 2016]. We present pseudocode for fitting a rank-one component with both exponential decay and underapproximation constraints in Algorithm 3. In this function, PAVAD_ECREASING_ implements a projection onto the set of exponentially decreasing vectors using the OASIS algorithm from [Friedrich et al., 2017].

##### Algorithm 1: Photocurrent Removal with Constraints (PHoRC)

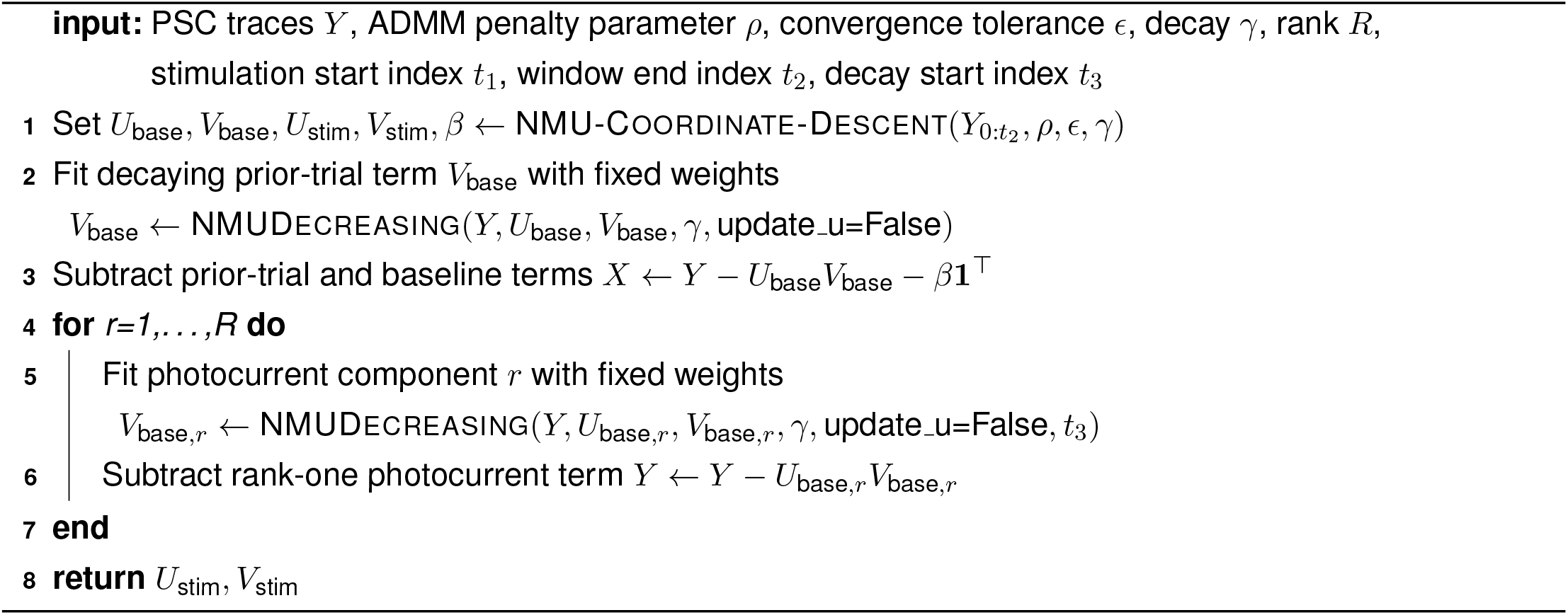

#### 4.3.5 Batched photocurrent subtraction

As shown in Figure 2, we observed slight power-dependent changes in photocurrent kinetics. We therefore adopted a batching strategy when estimating photocurrents, which we observed to improve performance on real data. To begin, we sort all photocurrents by the energy in the signal during stimulation. We then run the PhoRC algorithm (Algorithm 1) on batches of sorted traces. For all experiments shown, we used a batch size of 100 traces.

##### Algorithm 2: NMU-Coordinate-Descent

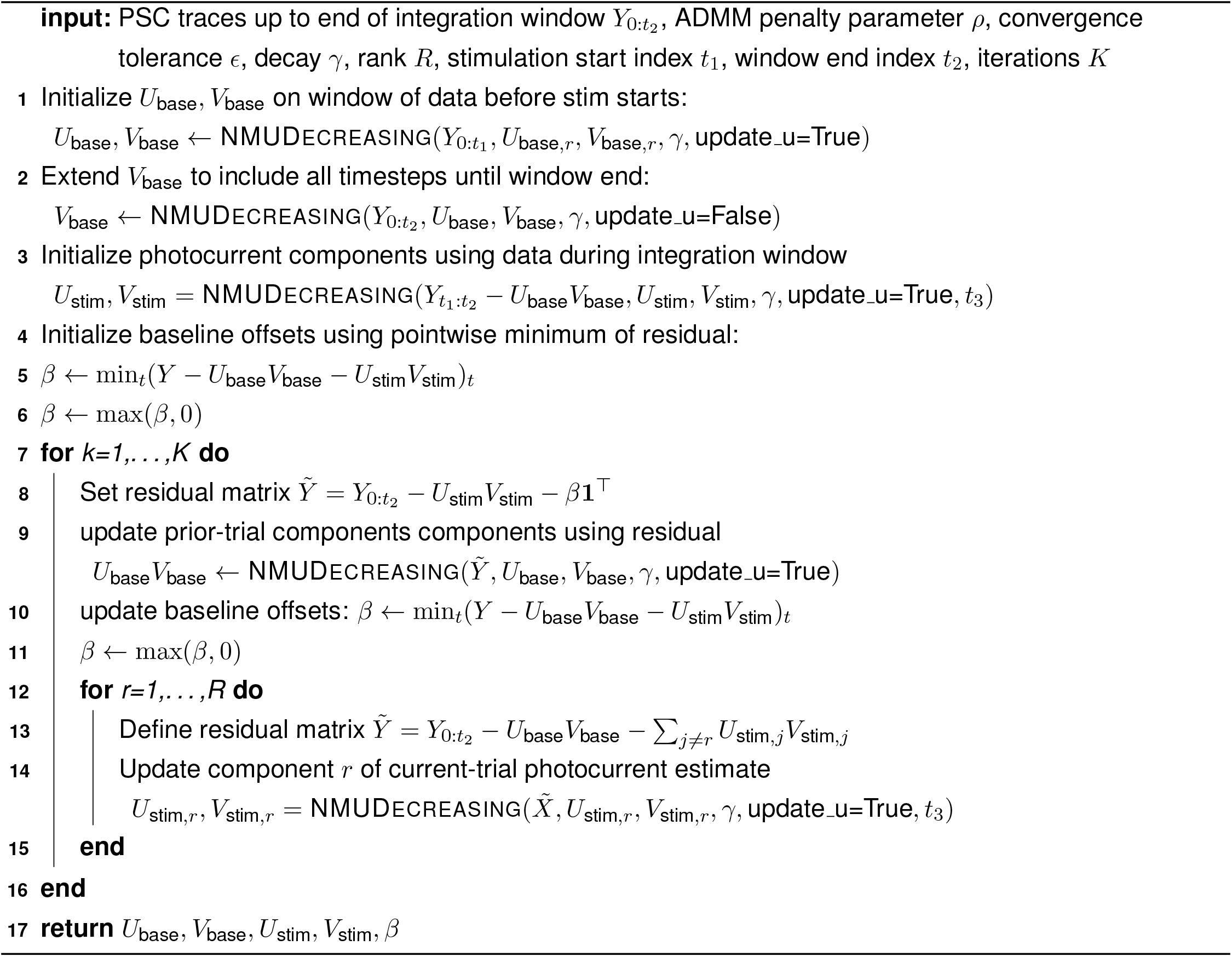

##### Algorithm 3: NMUDecreasing

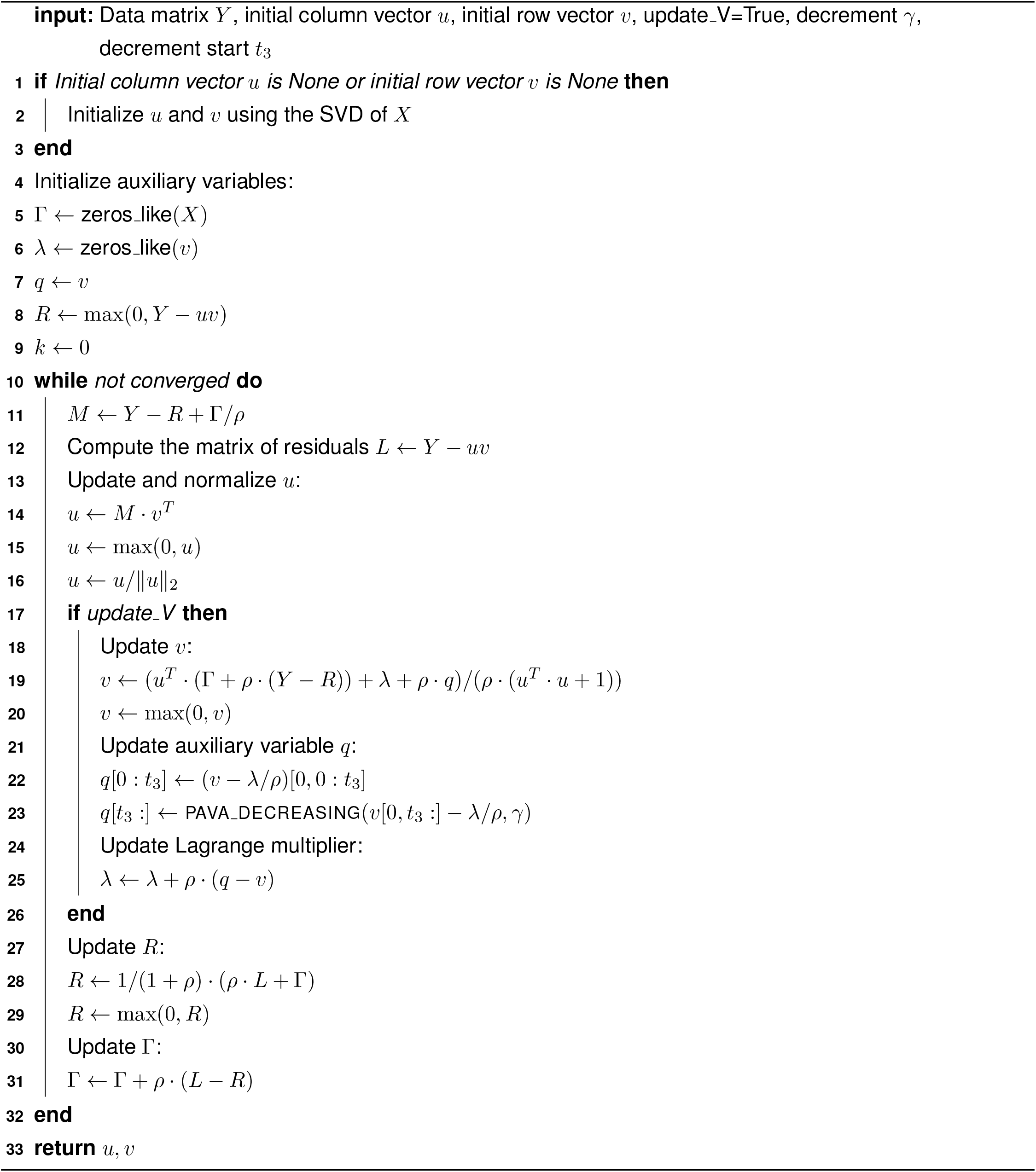

#### 4.3.6 Setting PhoRC hyperparameters

Two main hyperparameters determine how aggressive PHoRC is when subtracting photocurrents. These are the photocurrent integration window (see section 2.4.2) and the rank of the approximation (see subsection 4.3). Recall that PhoRC proceeds in two steps, first using a short window of data (the integration window) to learn the photocurrent weights *U* and subsequently estimating the waveform *V* with *U* held fixed. Longer integration windows result in more complete photocurrent removal, at the expense of occasionally subtracting low-latency PSCs. We found in simulation that a good starting point for the integration window is 3 ms, since this resulted in minimal subtraction of PSCs. However, in cases of extreme photocurrent contamination, we found that extending this window to the full duration of laser stimulation reduced residual photocurrent artifacts.

In our results, we used a 5 ms integration window on all grid datasets, and a 3 ms integration window on all targeted datasets, matching the respective pulse durations used in those experiments.

Including more components in the low-rank approximation also makes subtraction more complete, at the expense of sometimes partially subtracting PSCs. We found that when photocurrents were vastly larger than PSCs, a rank-two approximation worked best, possibly because there was more variability in the photocurrent waveform in those cases. In cases where photocurrents were around the same magnitude as PSCs, a rank-one approximation worked well. We used rank-two approximations for all of the grid experiments, and rank-one approximations for the targeted experiments.

### 4.4 Connectivity inference and estimation of putative connection waveforms

To assess the impact of photocurrents on connectivity, we relied on the CAVIaR pipeline from [Triplett et al., 2022]. This pipeline comprises two stages: neural waveform demixing (NWD), and connectivity inference. The NWD stage uses a neural network to denoise data and correct for confounded baselines when using short ISIs. This stage also subtracts any PSCs which are not initiated within a window characteristic of monosynaptic transmission, typically 3-12 ms. The connectivity inference stage uses a statistical model to infer which trials were successful in eliciting an upstream spike, and subsequently to learn an effective synaptic weight for each stimulation target. In all results, we used both stages of the pipeline when inferring connectivity (e.g Figs. 3, 7). In the presenence of photocurrent artifacts, we found that combining PhoRC with NWD yielded better results than applying either technique independently.

As described in [Triplett et al., 2022], the connectivity inference stage of CAVIaR allows for the computation of putative connection waveforms as follows. During connectivity inference, CAVIaR infers a matrix Λ, in which Λ_*ij*_ represents the posterior probability that cell *i* spiked on trial *j*. Using this presynaptic spike matrix **Λ**, we can obtain accurate PSC waveforms **r**_*n*_ ∈ ℝ^*T*^ using ridge regression. Collecting the waveforms in the rows of a matrix **R**, we obtain an estimate for **R** by solving the *L*_2_ problem

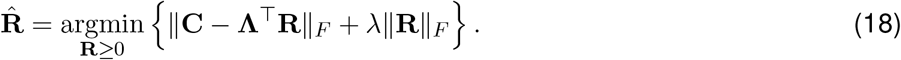

In all results showing connection waveforms, we computed this regression using only trials at the highest laser power.

### 4.5 Selecting representative traces

In order to demonstrate PhoRC’s performance, we found it useful to examine the model’s behavior on a small subset of traces collected during the experiment. In most cases, we extracted traces from two categories: those with the largest putative synaptic current components, and those with the the largest putative photocurrent components.

To extract traces with large synaptic currents, we first ran PhoRC to remove photocurrents, and then applied NWD to denoise and correct baselines. NWD is trained to isolate PSCs which onset in an “admissible window” around stimulation onset, and thus it tends to reduce synaptic currents caused by spontaneous activity [Triplett et al., 2022]. We thus treated traces with the largest NWD outputs as being likely to contain evoked synaptic currents.

To extract traces with large putative photocurrents, we simply used the magnitude of the PhoRC estimate, restricted to the time during stimulation. Thus, wherever we plotted representative traces (e.g Figure 3, Figure 7, and Supplementary Figures S1-S6), we included 10-15 traces with the largest NWD outputs, and 10-15 traces with the largest PhoRC outputs.

### 4.6 Validation with simulated connnectivity mapping experiments

For the simulated results in Figure 5, we simulated a continuous mapping experiment with a population of 100 presynaptic neurons, assigned at random to be connected to the postsynaptic cell with probability 0.1. Using the connectivity mapping simulator from [Triplett et al., 2022], we obtain simulated PSCs and electrical noise for each trial. In the interest of brevity, we refer the reader to [Triplett et al., 2022] for a detailed description of these connectivity mapping simulations. We note that these simulations assign a canonical PSC waveform to each connected cell, and that this waveform has power-dependent latency as we observed in the real data (see e.g Figure 1).

A fraction of cells were chosen with probability *p* (the “photocurrent fraction”) to evoke direct photocurrent in the postsynaptic cell. Concretely, let *q*_*n*_ = 1 if cell *n* evokes photocurrent; then

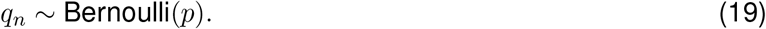

Conditional on *q*_*n*_ = 1, the maximal mean evoked photocurrent for presynaptic cell *n* was drawn from a gamma distribution:

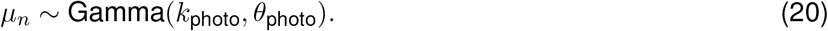

Finally, we draw the photocurrent amplitude contributed from presynaptic target *n* on trial *k* as

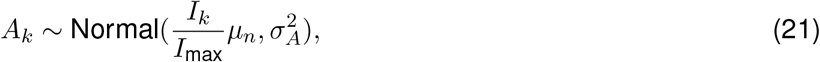

where *I*_max_ represents the maximum power used during the experiment, and *I*_*k*_ is the laser power on trial *k*. We can occasionally sample values for *A*_*k*_ which are below zero, so we simply take these to be zero. Thus *µ*_*n*_ represents the average evoked photocurrent amplitude when stimulating at the highest laser power. This model introduces variability, controlled by 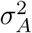, when stimulating the same presynaptic target, as we observe in the real data. The computed amplitude *A*_*k*_ is used to scale the photocurrent waveform, as we describe below.

In order to account for prior and next-trial effects, we simulated a continuous intracellular voltage recording, which was then broken down into overlapping 45ms snippets. This captures scenarios in which photocurrents and PSCs from the prior trial may not have decayed to zero before the next stimulus. Simulated photocurrents used a variant of the double two-state opsin model described by [Schoeters et al., 2021], represented by the following set of equations:

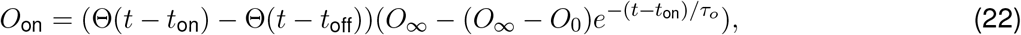

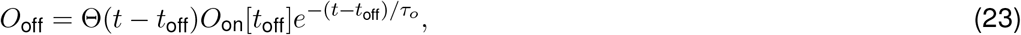

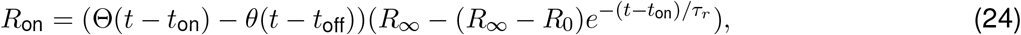

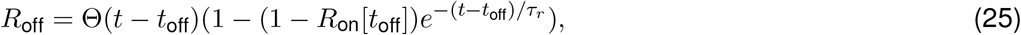

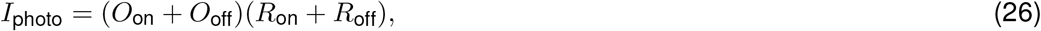

where Θ(*t*) is the heaviside step function, and *I*_photo_ is the induced photocurrent. The parameters governing the shape of the photocurrent waveform, *R*_inf_, *O*_inf_, *τ*_*o*_, *τ*_*r*_, were selected uniformly at random from a range of values to match variability in photocurrent kinetics in the real data (see Figure 2). For each simulated experiment, we re-sampled values of these parameters, giving a simulated postsynaptic cell its own photocurrent waveform. At each trial, we add scaled photocurrents *A*_*nk*_*I*_photo_ to simulated PSCs and noise. In our simulations, we use a 5ms pulse duration, and include 5 ms of context from the prior trial so that *t*_on_ = 5 ms and *t*_off_ = 10 ms.

### 4.7 Validation with hybrid data

For the hybrid data validation presented in Figure 6, we began with opsin-negative grid mapping datasets collected using the Ai203 mouse line [Bounds et al., 2021] and added synthetic photocurrents on top of recorded ground-truth traces. For these simulations, it was important that the photocurrent contamination had a spatial footprint, as we see in the real data. To accomplish this, we use a three dimensional Gaussian density to modulate the mean evoked photocurrent at each point in space. Let **x**_*k*_ be a point on a three-dimensional grid stimulated during trial *k*, let *I*_*k*_ be the power delivered at trial *k*, and let **c** be the location of the postsynaptic cell. We define the mean photocurrent amplitude at a given location and laser power to be

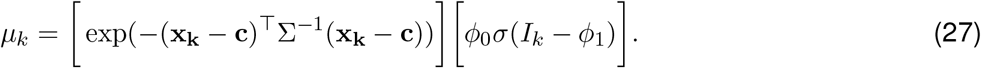

Here, *µ*_*k*_ is a mean used to sample the photocurrent amplitude, Σ is a diagonal matrix determining the spatial extent of the photocurrent response. *ϕ*_1_ and *ϕ*_0_ are parameters governing how *µ*_*k*_ changes with laser power, and *σ* is the sigmoid function. Thus, the first bracketed term scales *µ*_*k*_ with distance from the postsynaptic cell, and the second bracketed term scales *µ*_*k*_ with laser power. To introduce random variability when stimulating the same target location, we sample *A*_*k*_, the photocurrent amplitude on trial *k*, as

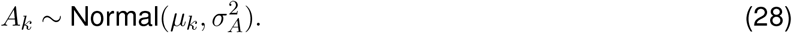

Using simulated photocurrents from Equation 26, we added these scaled photocurrents to the ground-truth grid maps. We then ran PHoRC and computed mean-squared error (MSE) by comparing the recovered PSC responses to the ground-truth data.

**Figure S1:**
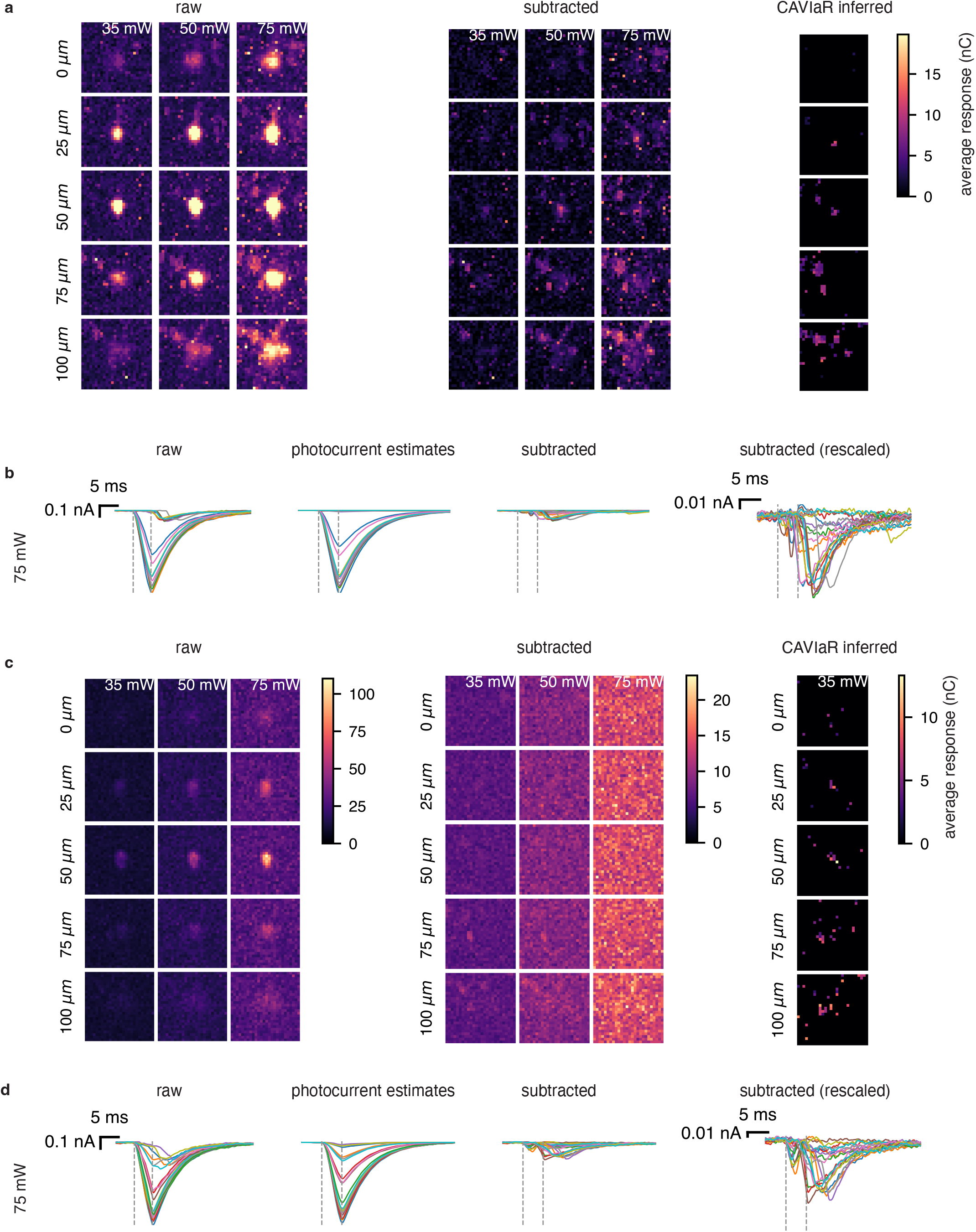
Cell 1 subtraction detail with comparison between single-target and compressed-sensing based mapping. Same dataset as shown in Figure 1, Figure 3, and Figure 7 **a**, Single-target mapping. Raw, estimated, and subtracted maps are shown for five planes (rows) and three powers (columns). The rightmost column shows the result of applying CAVIaR to estimate synaptic weights, which effectively merges data across laser powers. Patched cell is on plane 50 *µ*m. **b**, Raw, estimated photocurrents, and subtracted traces for highest laser power. Since the photocurrents and estimates are so much larger than synaptic currents, the rightmost column shows the subtracted traces rescaled for legibility. Traces were selected by finding the 15 largest estimated synaptic responses, and the 15 largest estimated photocurrents. **c**, Same as **a** for ensemble mapping with the same postsynaptic cell. For multispot data, raw and subtracted maps are created by averaging all ensembles in which a given voxel was stimulated, thus we expect these maps to appear “noisier” than in the single-target case. **d**, Same as **b** for ensemble mapping of the same postynaptic cell.

**Figure S2:**
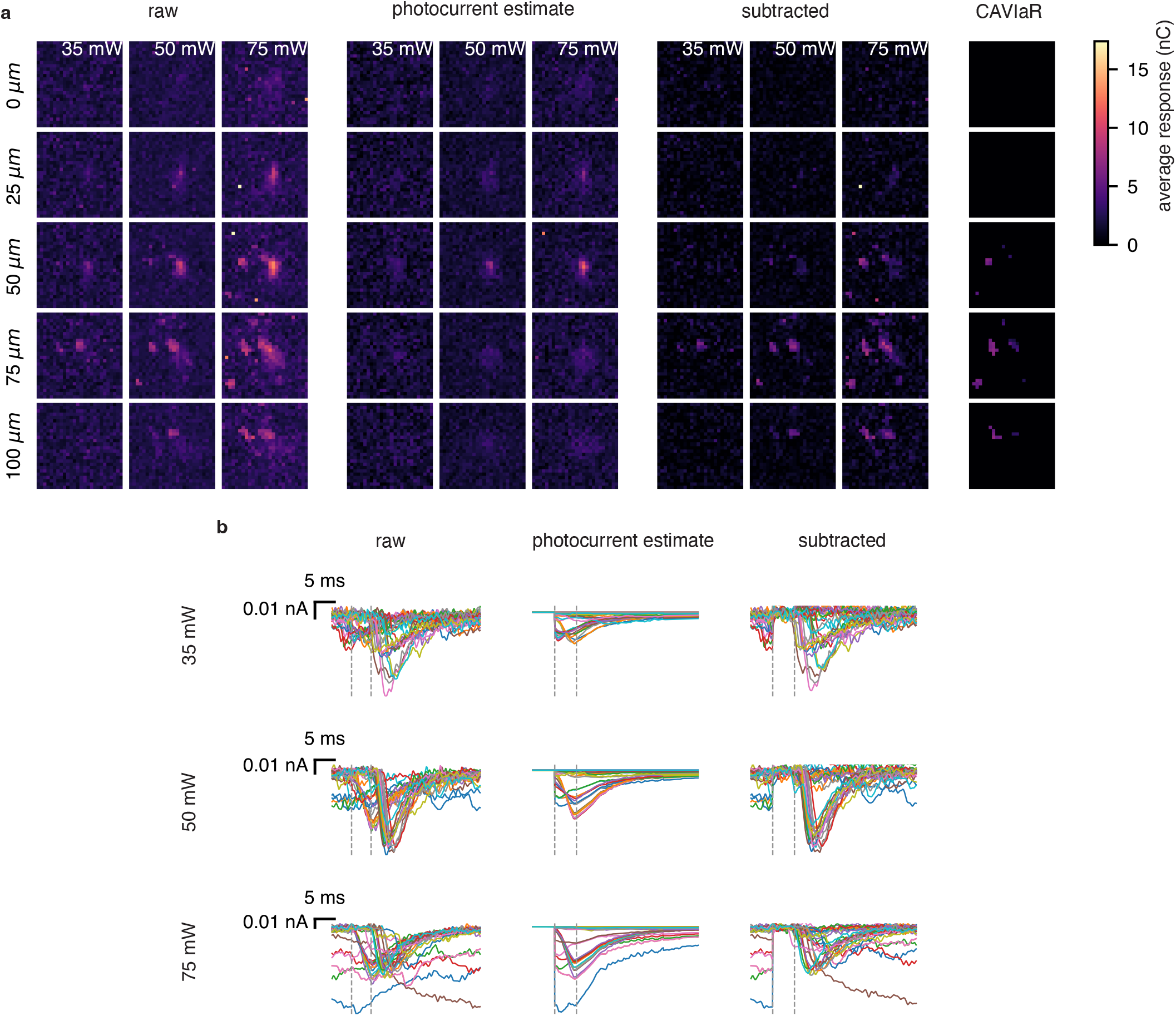
Cell 2 subtraction detail. **a** Full grid mapping dataset showing subtraction performance. Raw, estimated, and subtracted maps are shown for five planes (rows) and three powers (columns). For legibility, color-scale is truncated to match the range of the subtracted data. The rightmost column shows the result of applying CAVIaR to estimate synaptic weights, which effectively merges data across laser powers. Patched cell is on plane 50 *µ*m. **b** Raw, estimated photocurrents, and subtracted traces for three powers. Traces were selected by finding the 15 largest estimated synaptic responses, and the 15 largest estimated photocurrents. As seen in the bottom row, PhoRC occasionally subtracts traces with corrupted baselines.

**Figure S3:**
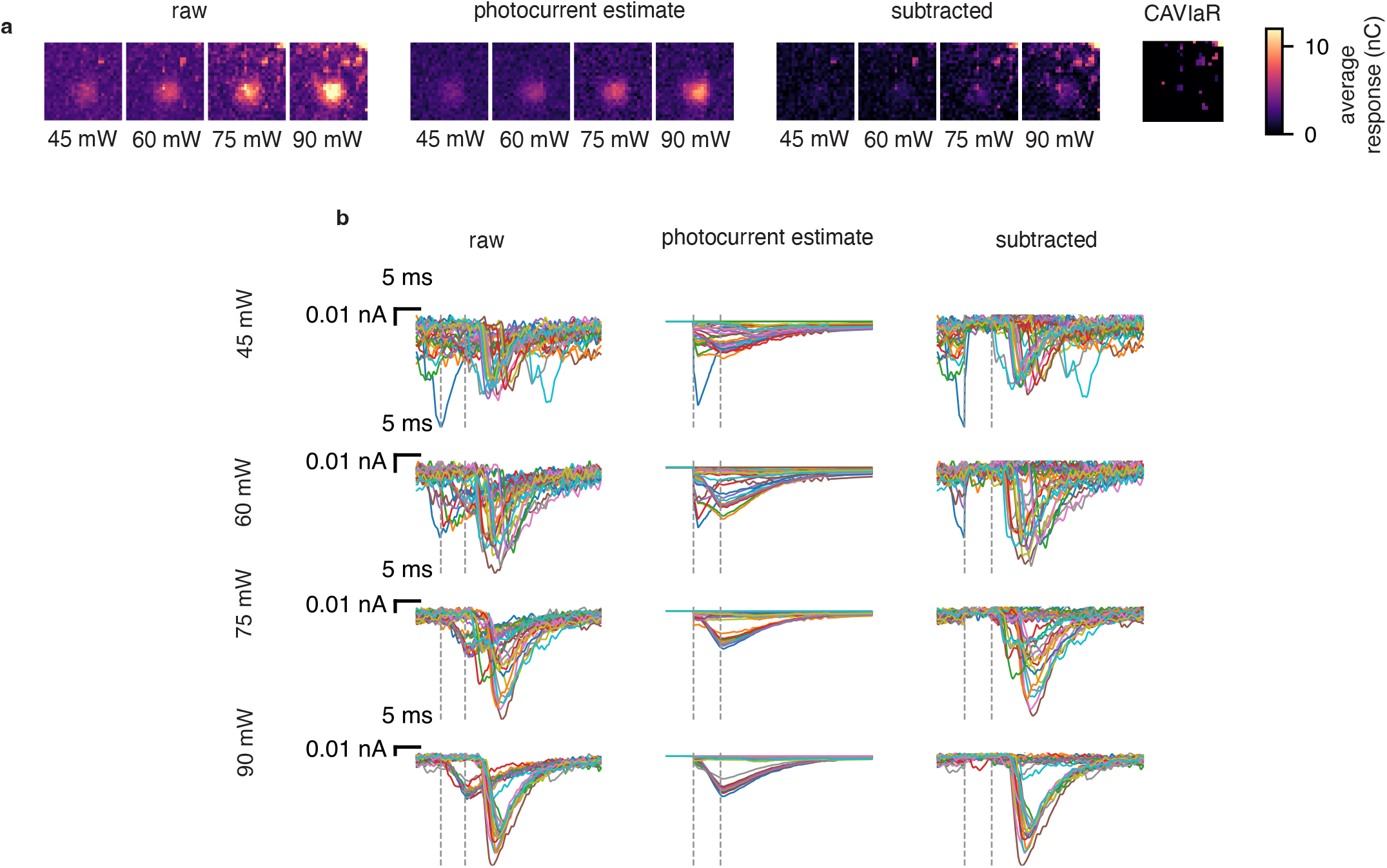
Cell 3 subtraction detail. **a** Full grid mapping dataset showing subtraction performance. Raw, estimated, and subtracted maps are shown for three powers (columns). For legibility, color-scale is truncated to match the range of the subtracted data. The rightmost column shows the result of applying CAVIaR to estimate synaptic weights, which effectively merges data across laser powers. Patched cell is on plane 50 *µ*m. **b** Raw, estimated photocurrents, and subtracted traces for three powers. Traces were selected by finding the 15 largest estimated synaptic responses, and the 15 largest estimated photocurrents. Note that in some cases PhoRC mistakenly captures very low-latency PSCs. This is expected behavior, since these PSCs occur concurrently with stimulation onset (and are therefore likely indicative of spontaneous activity rather than a monosynaptic connection).

**Figure S4:**
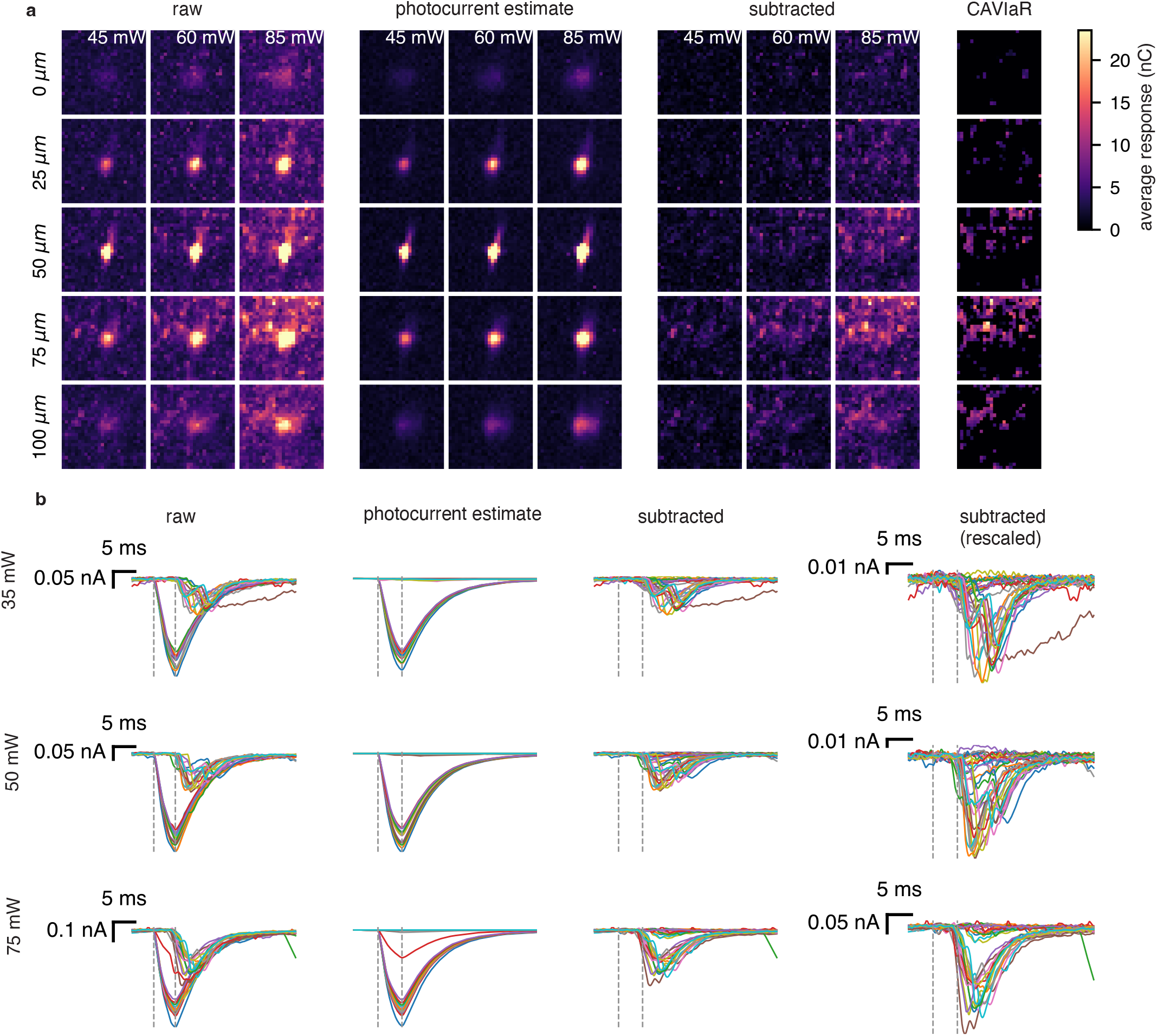
Cell 4 subtraction detail. **a** Full grid mapping dataset showing subtraction performance. Raw, estimated, and subtracted maps are shown for five planes (rows) and three powers (columns). For legibility, color-scale is truncated to match the range of the subtracted data. The rightmost column shows the result of applying CAVIaR to estimate synaptic weights, which effectively merges data across laser powers. Patched cell is on plane 50 *µ*m. **b** Raw, estimated photocurrents, and subtracted traces for three powers. Traces were selected by finding the 15 largest estimated synaptic responses, and the 15 largest estimated photocurrents.

**Figure S5:**
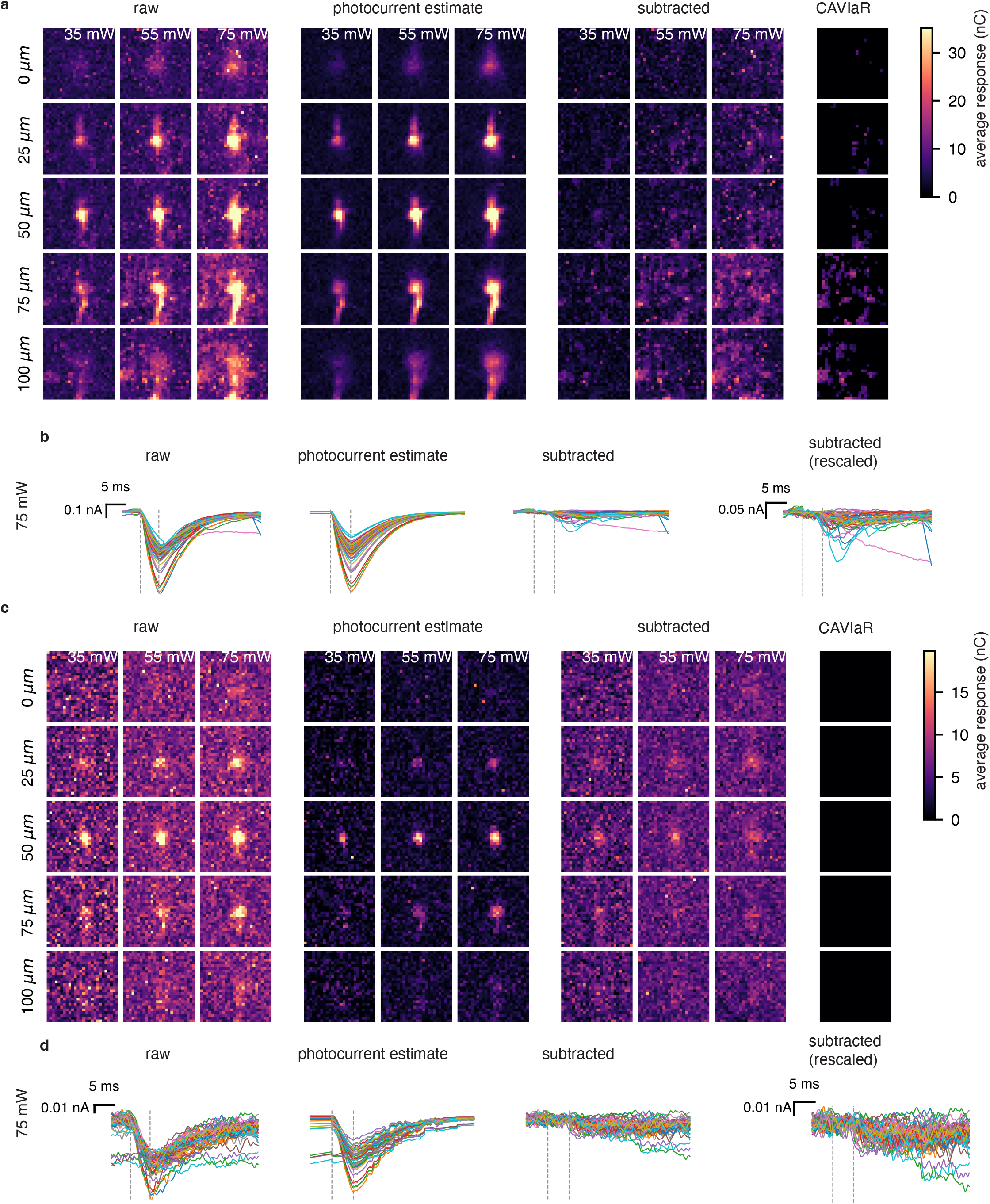
Cell 5 subtraction detail with synaptic transmission block. **a** Full grid mapping dataset during the control block. Raw, estimated, and subtracted maps are shown for five planes (rows) and three powers (columns). For legibility, color-scale is truncated to match the range of the subtracted data. Patched cell is on plane 50 *µ*m. **b** Raw, estimated photocurrents, and subtracted traces for the highest laser power. Traces were selected by taking the responses with the largest estimated photocurrent component. **c**, Same as **a** but during the NBQX block. **d**, Same as **b** but during the NBQX block. Traces were selected by taking the responses with the largest estimated photocurrent component.

**Figure S6:**
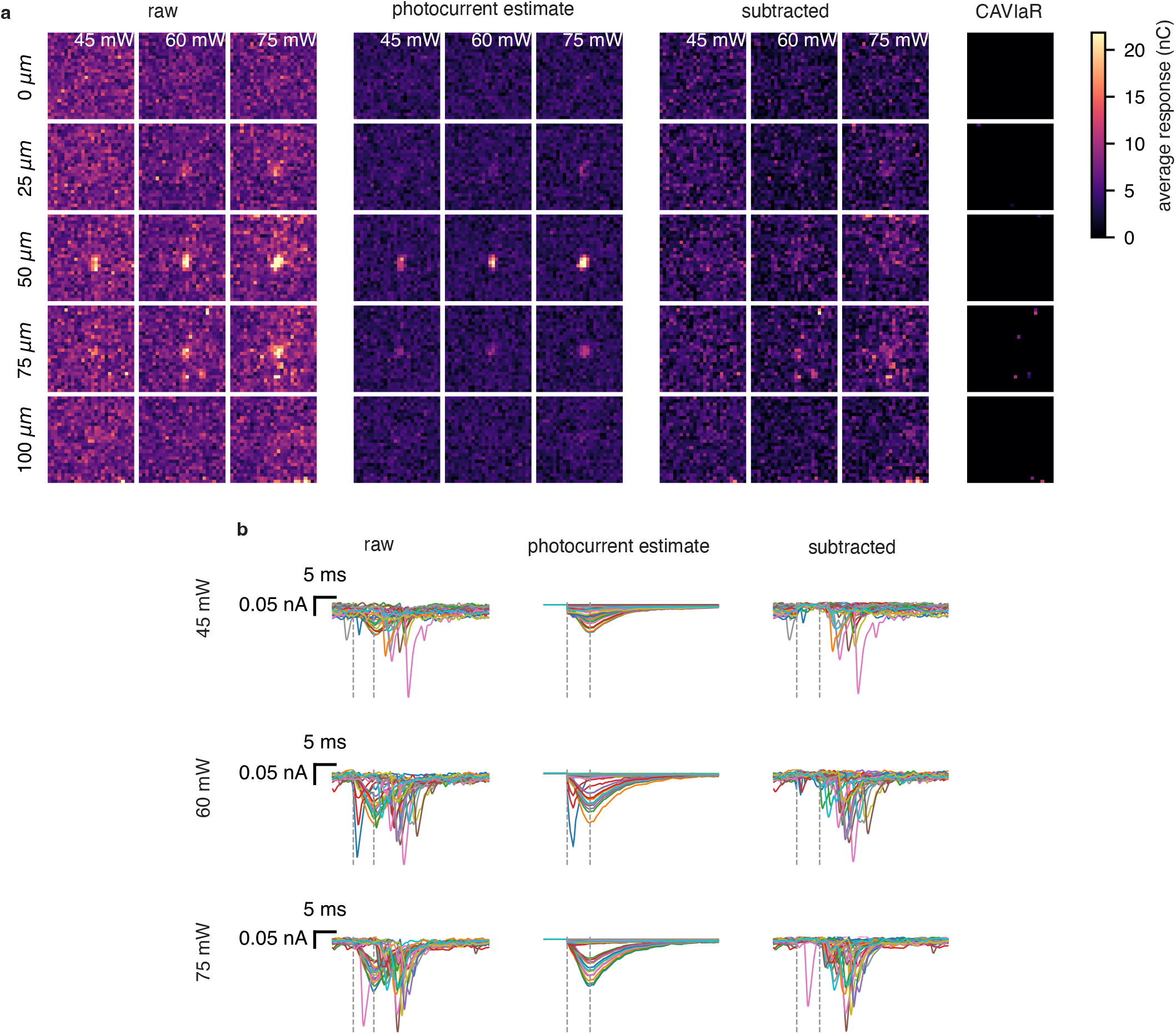
Photocurrent subtraction for pyramidal to PV mapping. This dataset represents a putative PV patch, recognizable by the very sharply decaying EPSCs. **a** Full grid mapping dataset showing subtraction performance. Raw, estimated, and subtracted maps are shown for five planes (rows) and three powers (columns). For legibility, color-scale is truncated to match the range of the subtracted data. The rightmost column shows the result of applying CAVIaR to estimate synaptic weights, which effectively merges data across laser powers. Patched cell is on plane 50 *µ*m. **b** Raw, estimated photocurrents, and subtracted traces for three powers. Traces were selected by finding the 15 largest estimated synaptic responses, and the 15 largest estimated photocurrents. As seen in the middle row, PhoRC occasionally subtracts PSCs which occur around the time of stim onset. This is to be expected, and their very low latencies suggest that these PSCs are likely due to spontaneous activity.

**Figure S7:**
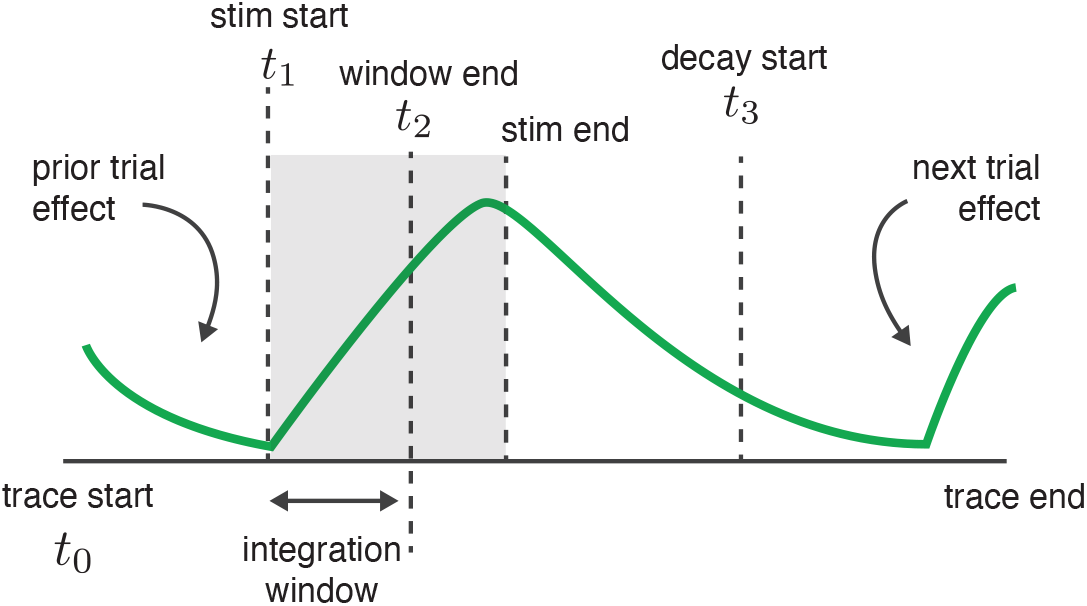
Accounting for prior and subsequent trial effects. The integration window begins at the time of laser onset (*t*_1_) and ends at a user-defined time (*t*_2_), typically chosen to be 3-5ms after laser onset. To avoid including photocurrents or synaptic currents from the subsequent trial, we enforce an exponential decay constraint beginning at time *t*_3_.

